# Intelligence and Visual Mismatch Negativity: Is Pre-Attentive Visual Discrimination Related to General Cognitive Ability?

**DOI:** 10.1101/2022.03.01.482097

**Authors:** Kirsten Hilger, Matthew J. Euler

## Abstract

Electroencephalography (EEG) has been used for decades to identify neurocognitive processes related to intelligence. Evidence is accumulating for associations with neural markers of higher-order cognitive processes (e.g., working memory); however, whether associations are specific to complex processes or also relate to earlier processing stages remains unclear. Addressing these issues has implications for improving our understanding of intelligence and its neural correlates. The mismatch negativity (MMN) is an event-related brain potential (ERP) that is elicited when, within a series of frequent standard stimuli, rare deviant stimuli are presented. As stimuli are typically presented outside the focus of attention, the MMN is suggested to capture automatic pre-attentive discrimination processes. However, the MMN and its relation to intelligence has largely only been studied in the auditory domain, thus preventing conclusions about the involvement of automatic discrimination processes in humans’ dominant sensory modality vision. Electroencephalography was recorded from 50 healthy participants during a passive visual oddball task that presented simple sequence violations as well as deviations within a more complex hidden pattern. Signed area amplitudes and fractional area latencies of the visual mismatch negativity (vMMN) were calculated with and without Laplacian transformation. Correlations between vMMN and intelligence (Raven’s Advanced Progressive Matrices) were of negligible to small effect sizes, differed critically between measurement approaches, and Bayes Factors provided anecdotal to substantial evidence for the absence of an association. We discuss differences between the auditory and visual MMN, the implications of different measurement approaches, and offer recommendations for further research in this evolving field.

**HIGHLIGHTS:**

- Testing whether intelligence is related to automatic visual discrimination
- Visual mismatch negativity (vMMN) as a neural indicator of pre-attentive processing
- No association between intelligence and vMMN amplitudes or latencies
- Critical differences between auditory and visual MMN?
- Results partly depend on different measurement approaches

## Introduction

Intelligence is a psychological construct that includes the ability to understand complex ideas, to adapt effectively to the environment, to learn from experience, and to engage in various forms of reasoning (Neisser et al., 1996). Scientific tests have been developed that allow the calculation of so-called intelligence quotients. Respective scores try to capture the general cognitive ability level of a person and have been shown to predict educational and occupational success (Schmidt & Hunter, 2004) as well as positive life outcomes such as health and longevity (Deary et al., 2004). Understanding the cognitive processes contributing to different levels of intelligence is therefore an important aim of ongoing research across multiple scientific disciplines.

The study of individual variation in neural parameters assessed during specific tasks (involving specific cognitive processes) provides a means to gain insight into the question of how certain processes may contribute to individual differences in intelligence. Neuroimaging research has identified neural correlates of intelligence in brain structure (e.g., Haier et al., 2004; Hilger et al., 2020a), brain function (e.g., Gray et al., 2003; Lipp et al., 2012), and in intrinsic brain connectivity (e.g., Hilger et al., 2020b; for general reviews on neuroimaging correlates of intelligence see Basten et al. 2015; Hilger & Sporns, 2021; Jung and Haier 2007). However, such research often provides only limited insights into intelligence-critical *processes*, as the identified neural parameters that covary with differences in intelligence (e.g., intrinsic connectivity in region of the dorsal attention network, DAN) were typically not assessed during an intelligence-critical cognitive processes. Rather they were assessed during the resting state or during tasks not necessarily related to intelligence (e.g., movie watching; Haier et al., 2003). Although findings from previous investigations allow for vague interpretations about the meaning of these identified intelligence-related parameters (e.g., other studies that showed that regions of the DAN were associated with attentional processes), any such inferences are rather speculative, especially because most neural parameters have been associated not only with one but with many different processes (reverse inference problem, Nathan & Del Pinal, 2017; Poldrack, 2008, 2011, 2015).

The study of event-related brain potentials (ERP) with electroencephalographic recordings during specific tasks allows to more directly study the relation between variability in intelligence and its association with process parameters. Significant associations between intelligence and neural correlates of higher-order cognitive processes have been reported frequently, such as working memory (e.g., Stipacek et al., 2003) or attentional control (e.g., Schubert et al., 2017, 2022; for review see Hilger et al., 2022). However, the relevance of lower-order processes for intelligence is unclear. Investigating this issue is important for clarifying the nature of intelligence itself, as a construct that is either exclusively concerned with higher-order capacities, or which (for example) may reflect aggregated variation across a hierarchy of lower-order and other processes (Euler, 2018). Support for this idea comes from research showing small but reliable correlations between intelligence and apparently ‘basic’ processes, such as simple reaction time (Deary et al., 2001) and pitch or color discrimination (Acton & Schroder, 2001), and in the neural domain from studies showing associations between intelligence and non-task-related early visual processing (Euler et al., 2015), and even pre-stimulus anticipatory potentials (McKinney & Euler, 2019). Stated another way, while it is clearly important to assess relations between intelligence and the neural processes that govern higher-order functions, attaining a complete understanding of intelligence equally requires investigating processes near the bottom of the cognitive hierarchy. As such, it has not yet been sufficiently clarified whether intelligence-related associations are specific to complex higher-order cognitive processes or whether there also exist associations at early (pre-attentive) processing stages.

The auditory mismatch-negativity (MMN) has evolved as promising marker of individual differences in intelligence (e.g., De Pascalis et al., 2014; De Pascalis & Varriale, 2012; Houlihan & Stelmack, 2012; Sculthorpe et al., 2009; Troche et al., 2009, 2010). In general, the MMN is elicited when, within a series of frequent standard stimuli, rare deviant stimuli are presented (e.g., tones of higher pitch). As stimuli are typically presented outside the focus of attention, the MMN is suggested to capture automatic pre-attentive discrimination processes (Näätänen et al., 2007). Associations between higher intelligence and larger (i.e., more negative) MMN amplitudes have been observed in multiple studies (De Pascalis et al., 2014; De Pascalis & Varriale, 2012; Houlihan & Stelmack, 2012; Sculthorpe et al., 2009; Troche et al., 2009, 2010) as well as a relation between higher intelligence and shorter MMN latencies (De Pascalis et al., 2014; De Pascalis & Varriale, 2012; Sculthorpe et al., 2009). However, associations were mostly of moderate effect size (∼ *r* = - .15 to *r* = -.42) and different studies also failed to find respective relations (e.g., Bauchamp & Stelmack, 2006; De Pascalis & Varriale, 2012; Troche et al., 2010), suggesting a more heterogeneous picture.

Although vision constitutes humans’ dominant sensory modality (e.g., Hutmacher, 2019), automatic discrimination processes via the MMN have typically been investigated in the auditory domain only. Experimental paradigms for assessing the MMN also in the visual domain were first developed in 1990 (Cammann, 1990; Czigler & Csibra, 1990; Nyman et al., 1990), and the available literature suggests that violations in both simple and more complex visual patterns that are presented outside the focus of attention elicit an electrophysiological component similar to the auditory MMN, i.e., the visual MMN (vMMN; e.g., Stefanics et al., 2011, 2014; Zeng et al., 2022; for review see Pazo-Alvarez et al., 2003). To our knowledge, just one study has examined an association between vMMN and intelligence (Liu et al., 2015). Specifically, Lui et al. (2015) compared groups of average and high IQ adolescents during completion of two emotional vMMN conditions: A happy condition in which neutral faces represented standard stimuli and happy faces were deviants, and a fearful condition in which neutral faces served again as standards and fearful facial expressions as deviants. All stimuli were presented outside the focus of attention, i.e., in addition to a target change detection task. Results indicated that in the early vMMN window (50-130 ms), the high IQ group showed larger vMMN amplitudes in the happy condition, with no differences for the fearful condition. For the later vMMN window (320-450 ms), the high IQ group expressed larger vMMNs over frontal and occipito-temporal areas in the fearful condition, while the average IQ group had larger vMMN amplitudes over occipito-temporal areas in the happy condition. However, as this study investigated an emotional vMMN elicited by rather complex perceptual stimuli (facial expressions), it is unclear how the findings relate to auditory MMN research where the MMN is typically elicited by unemotional and overall simple perceptual discrimination processes. Whether variations in the vMMN indexing simple visual discrimination processes relate to individual differences in intelligence therefore still constitutes an open question.

Here, we address this gap and transfer an established experimental vMMN paradigm (Stefanics et al., 2011) to research in the field of intelligence. We present two different ways to compute the vMMN as elicited by simple and more complex rule violations and use state-of-the-art operationalizations of its amplitude and latency. To preview our results, we observed a clear vMMN, replicating previous work. However, neither its amplitude nor its latency was related to variation in intelligence. Bayes Factors provided anecdotal to substantial evidence for the absence of an association, depending on the specific measurement approach. We conclude with a discussion of limitations and recommendations for further research in this evolving field.

## Method

### Participants and Assessment of Intelligence

60 right-handed students from Goethe University Frankfurt completed the experiment for monetary compensation or student credits. The size of this sample was determined by an a priori power calculation in combination with monetary feasibility. Specifically, based on previous work on the auditory MMN and intelligence (e.g., Sculthorpe et al., 2009), we expected an effect size ∼.35, which resulted, when ensuring 80% statistical power, in a required sample size of *N* > 49 participants (G*Power, Faul et al., 2007; two-tailed, α = .05). As we expected that some participants need to be excluded, we collected data of 60 persons. Students with a Major or Minor study subject in Psychology were excluded. All participants had self-reported normal or corrected-to-normal visual acuity and no history of psychiatric or neurological diseases. The procedures were approved by the local ethics committee (# 2015-201) and informed written consent according to the Declaration of Helsinki was obtained from all participants. Seven participants completed an earlier version of the protocol and were excluded due to insufficient numbers of trials. One additional participant was excluded due to EEG acquisition failure, and two participants were excluded due to an insufficient number of useable trials after artifact correction (i.e., fewer than *n* = 40 artifact-free trials in any single condition), leaving a final sample of *N* = 50 subjects (13 men, 37 women). Demographic information for the final study sample is listed in Table 1. Intelligence was assessed in time-limited (40 min) group settings (10-12 participants) with Raven’s Advanced Progressive Matrices Set II (RAPM, Raven et al., 1998). The RAPM sum scores were used in all analyses as the primary variable of interest and varied in the final sample between 12 and 34 (*M* = 24.56, *SD* = 4.74; Table 1) corresponding to an IQ score between 65 and 123 (*M* = 92.32, *SD* = 14.97).

**Table 1.** Descriptive statistics for individual difference variables

|  | *M (SD)* | *Minimum* | *Maximum* |
| --- | --- | --- | --- |
| Age | 23.64 (3.62) | 18 | 33 |
| RAPM | 24.56 (4.74) | 12 | 34 |
| Reaction Time | 0.53 (0.04) | 0.43 | 0.66 |
| Accuracy | 0.97 (0.07) | 0.53 | 1.0 |

*Note:* $N = 50$ (13 men, 37 women). Reaction time (in seconds) and accuracy averaged across all task blocks. RAPM = Ravens Advanced Progressive Matrices sum score.

### Stimuli and Procedure

The paradigm used in the present study was adapted from Stefanics et al. (2011). The task was presented on a Windows computer and the distance between participant and screen (21.5-inch diagonal) was 120 cm. Experimental stimuli consisted of 24 red or green circles (diameter 2.5 cm; vertical distance between centers of stimuli: 4.5 cm; horizontal distance between centers of stimuli: 4 cm; total area covered by colored discs: 18.5 × 20.5 cm) and were shown for 100 ms on a computer screen with black background (see Figure 1). All stimuli were presented pairwise, defined by shorter (within-pair; 300 ms) and longer (between-pair; 800 ms) inter stimulus intervals (ISIs; full trial duration 1,300 ms).

**Figure 1.**
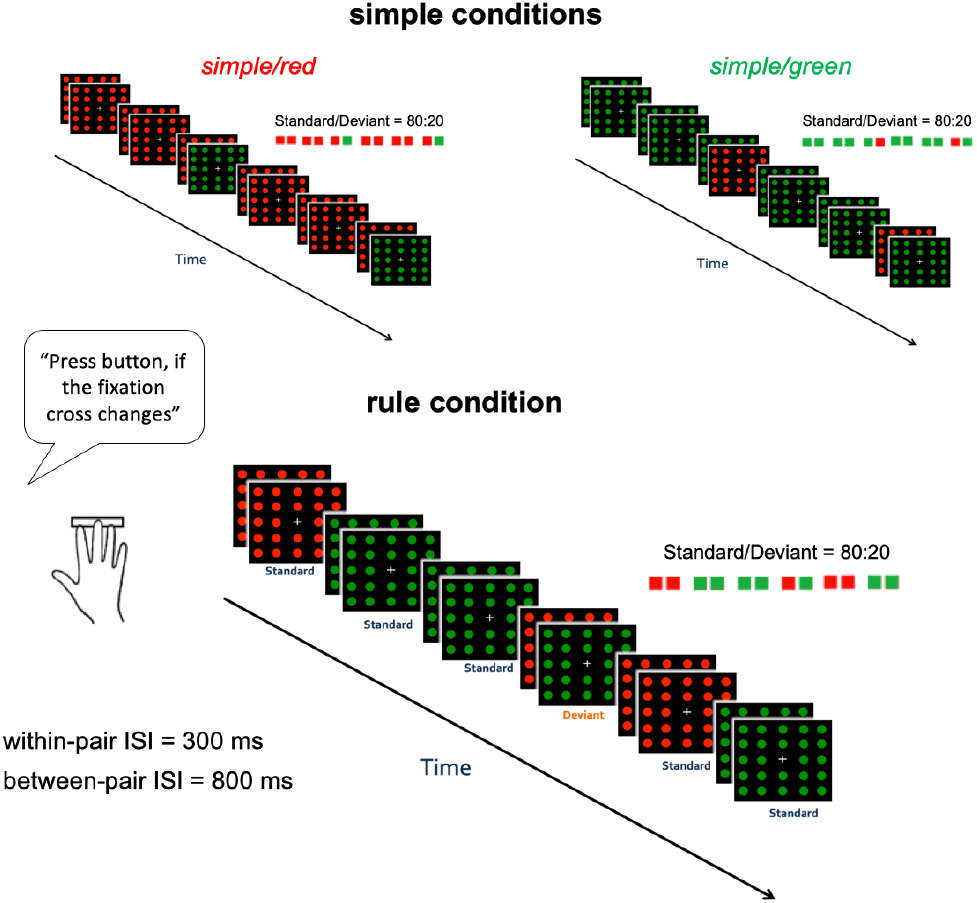
Schematic illustration of the passive oddball task. Top: Simple conditions (*simple/red* and *simple/green*). Bottom: Rule condition (*rule*). Stimuli were presented pairwise each for 100 ms with an ISI of 300 ms between two stimuli of the same pair and an ISI of 800 ms between two stimuli of different pairs (total trial duration: 1300 ms). Participants were instructed to focus on the fixation cross and to indicate via button press when they detect slight changes in the arm lengths of the fixation cross (occurring without any regularity on average 10 times per minute). Each block comprised 100 trials (100 pairs, 200 stimuli). The whole experiment consisted of 12 blocks, i.e., four blocks of each condition.

To study discrimination processes outside the focus of attention (passive oddball task), the participants were instructed to focus exclusively on a target, which consisted of a white fixation cross in the center of the screen. Participants were asked to indicate any change in the lengths of the arms of the cross by a speeded button press. Targets changed such that on half of the target trials, the horizontal arms of the cross were slightly elongated from 1.25 to 1.75 cm, while on the other half of target trials, the vertical arms were slightly elongated from 1 to 1.5 cm. Changes to both horizontal and vertical targets both corresponded to a ∼ 0.24° change in visual angle, and the relevant arms of the cross were always elongated symmetrically. The target presentation frequency was pseudo-randomized across trials, to ensure that targets were equally distributed across deviant and standard trials as well as across red and green stimuli. On average, targets (cross change) were presented 10 times per minute. The correct detection of a target as indexed by a button press within 800 ms after target onset was defined as hit, no reaction within 800 ms was registered as miss, and all other responses counted as false alarms.

The experiment consisted of three different conditions (see Figure 1). In the first oddball condition (*simple/red*) red stimuli served as standards and green stimuli served as deviants and were presented with a ratio of 80% to 20%, respectively. A reverse oddball condition had identical parameters, with the exception that green stimuli served as standards and red stimuli served as deviants (*simple/green*). The order of red and green stimuli within each block was pseudo-randomized to ensure that each block began with at least four standard stimuli. Further, two deviants were not allowed to be presented directly after one another but could occur at the first or the second position within a stimulus pair. In Figure 1, this is shown for simple/green, but not for simple/red. With respect to the attention control task, we ensured that a cross change could happen only once per stimulus pair (either at the first or second stimulus).

Finally, a more abstract, rule-based oddball task (*rule*) was implemented, where a standard was defined as a pair of two same-colored stimuli (red-red or green-green). The deviant in this condition was defined as a stimulus *pair* consisting of two different colors. Deviants were presented in 20% of trials. Note that as the decision about whether a stimulus pair represents a deviant or not always requires knowledge about the color of the second stimulus within a pair, deviance is in this condition always defined (and the corresponding neural mismatch response is initiated) by the occurrence of the second stimulus (see also Figure 1). The probability of red and green stimuli was equal within the whole block and the order of stimuli was pseudo-randomized such that at least four standard stimulus *pairs* were presented at the beginning of each block. Two deviant-pairs were not allowed to be presented after each other and cross changes could happen only once within a pair and not in two consecutively presented stimulus pairs. In the following, the first two oddball conditions (*simple/red* and *simple/green*) are labeled as ‘simple’ conditions, whereas the last condition is labeled as ‘rule’.

A total of 12 blocks was presented, with 4 blocks for each condition. For each subject, the order in which the 12 different pseudo-randomized stimulus blocks were presented within the whole experiment was generated purely by chance (completely randomized). Each block comprised 100 trials (i.e., 100 pairs, 200 stimuli) and took two minutes and 16 seconds. At the end of each block, the participants could decide to make a short break and were instructed to start the new block on their own when they felt ready.

### EEG Recording and Preprocessing

EEG data were recorded with 64 active Ag/AgCl electrodes (arranged in an extended 10–20 layout), using actiChamp amplifier (Brain Products GmbH, Gilching, Germany). FCz served as online reference, and AFz as the ground. The sampling rate was 1,000 Hz, impedance levels were kept below 10 kOm, and a low pass filter of 280 Hz was applied during acquisition (notch filter off). Two electrodes were placed below the left (SO1) and the right (SO2) eye to record ocular artifacts, and mastoid electrodes were placed behind both ears (M1, M2). Preprocessing and further analyses of EEG data were conducted in EEGLAB (Delorme & Makeig, 2004) and ERPLAB (Lopez-Calderon & Luck, 2014), and followed established standards for ERP processing and analysis (https://erpinfo.org/order-of-steps; Luck, 2014).

First, the continuous data were loaded into EEGLAB and a high-pass filter at 0.1 Hz was applied to correct any low-frequency drift. Data segments corresponding to task breaks were removed (time segments of ≥ 2000 ms or longer where no event code/trigger occurred, with a 500 ms buffer around the immediately preceding and following event code). The data were re-referenced to the average of all scalp electrodes, and two virtual bipolar EOG electrodes were created (consisting of Fp1 and SO1, Fp2 and SO2) to facilitate the identification of blink-related artifacts. The data were then epoched from −100 to 400 ms relative to stimulus onset (for each stimulus within a pair) and each epoch was linearly-detrended. This produced twelve sets of data epochs corresponding to the following 12 sub-conditions: standard red and deviant green stimuli in the first vs. second position from the simple/red condition (4 epoch types), standard green and deviant red stimuli in the first vs. second stimulus position from the simple/green condition (4 epoch types), and standard or deviant red and green stimuli from the rule condition (4 epoch types). To avoid any possible confounding of the ERP effects due to motor preparation, all stimuli presented within 800 ms after a fixation cross change were excluded from analyses. For the same reason, we excluded all trials in which a button was pressed although not required (false alarm).

The epoched data were then used to identify ‘bad’ electrodes, i.e., electrodes for which more than 4 % of trials would have been rejected due to non-blink-related artifacts, such as drift, or high-frequency noise by the automatic EEGLAB artifact-rejection procedure (voltage changes ≥ 60 µV on either side of a 100 ms window across 50 ms steps or voltage differences exceeding 100 µV within a 100 ms window, moving in 50 ms steps). The continuous (non-epoched) data of these electrodes were then manually reviewed and replaced via spherical-spline interpolation. Of note, our strict interpolation strategy resulted in, on average, 8.76 interpolated electrodes per subject (*SD* = 6.71; range: 0-26). However, we selected this very conservative artifact rejection approach to gain a very clean grand-average picture unaffected by idiosyncratic bad electrodes across participants. Importantly, of the nine electrodes used for statistical analyses (see below), very few were ultimately interpolated (*M* = 0.54 per subject; *SD* = .97; range: 0-3) suggesting minimal influences of our strict interpolation strategy on the vMMN estimation.

Thereafter, the same automated threshold-based routines were applied to now identify artifactual *epochs* containing either step-like artifacts in the ocular channels (those with voltage changes ≥ 60 µV on either side of a 100 ms window across 50 ms steps), or with voltage differences exceeding 100 µV within a 100 ms window, moving in 50 ms steps, across any scalp channels. All final data files were then manually reviewed again to ensure appropriate inclusion and exclusion of valid and artifactual trials by these algorithms. After rejecting trials contaminated due to artifacts, the remaining trials were used for calculating subject-specific ERPs. No participant had fewer than 43% of the total possible trials retained (minimum absolute number = 14/32 trials for the deviant rule conditions), with up to 100% of trials being retained in many participants and conditions.

### Event-Related Potentials

For the primary analyses, the cleaned epochs were then baseline-corrected relative to the 100 ms preceding the stimulus, and event-related potentials (ERPs) were created by averaging all epochs within each of the 12 epoch types. The resulting averaged ERPs were low-pass filtered at 10 Hz, to minimize high-frequency noise that can contaminate ERP latency measurements (Luck, 2014). Note that the analyses of within-subject condition effects focus on these epoch type-averaged ERPs (i.e., trial-averaged ERPs for each of the 12 epochs), where our aim is to establish the presence of a vMMN and its sensitivity to condition effects. In contrast, the individual differences analyses focus on ERP difference waves, which were derived by subtracting the ERP elicited by standard stimuli from the ERP elicited by deviant stimuli. For the simple task conditions, these difference ERPs were calculated based on standard and deviant stimuli within the same block of trials, and from the same position in the stimulus pair (1^st^ vs. 2^nd^ stimulus). This resulted in two difference waves created by subtracting red standards from green deviants in the first and second stimulus positions from the simple/red block (Green Odd 1 and Green Odd 2, where the designation “Odd” always refers to the color of the deviant stimulus), and two difference waves created by subtracting green standards from red deviants in the first and second stimulus positions from the simple/green block (Red Odd 1 and Red Odd 2). For the rule conditions, difference waves were created by subtracting red or green standards from their deviant counterparts in the opposite color (always in the second position; Red Odd Rule and Green Odd Rule). This process resulted in a total of six ERP difference waves that served as basis to identify the vMMNs.

#### ERP Measurement Strategy

As in most neuroscientific research, the calculation of ERPs implies many researchers’ degrees of freedom (Open Science Collaboration, 2015; Wicherts, et al., 2016), and it is still a matter of debate what the best analysis strategy actually is. Therefore, to increase the robustness of our findings against different choices in processing parameters we examined two different EEG referencing schemes:

##### A)The scalp-average referenced approach - direct replication of Stefanics et al. (2011)

For each subject-specific, trial-averaged ERP and ERP difference wave (vMMN), we used ERPLAB algorithms to calculate amplitudes and latencies (see next paragraph) in the period from 100-400 ms post-stimulus, at nine posterior electrodes (P3, Pz, P4, PO3, POz, PO4, O1, Oz, O2). These individual amplitude and latency values were then averaged across red and green stimulus conditions within each block (i.e., the two simple and one rule; matched by deviance and position) and used as input for the analyses of condition effects, with the goal of replicating the methods and findings of Stefanics et al. (2011). For the subsequent individual difference analyses, the relevant amplitude or latency values were averaged over all nine electrodes to produce a single value for each subject.

##### B)The Laplacian-transformed approach

The Laplacian transformation (also termed surface Laplacian [SL], Laplacian, scalp current density [SCD], or current source density [CSD]) is a mathematical simplification of Poisson’s equation (see e.g., Carvalhaes & de Barros, 2015) that relates current sources generated within the brain to the macroscopic potentials observable at scalp (i.e., the biophysical principle of volume conduction). The resulting transformation of the scalp topography produces a current density with a spatial extent (measured in µV/cm^2^) that emphasizes radially-oriented, superficial cortical sources, to the exclusion of deep focal sources or superficial but spatially-diffuse cortical activity (Nunez & Srinivasan, 2006). In contrast to the “traditional” analysis approach, the Laplacian transformation has two major advantages: A) It is free of any reference and B) the resulting signal is closer to the dominant cortical source of the signal, i.e., less confounded by volume conduction (for an in-depth review and discussion see Kayser & Tenke, 2015). Thus, by contrasting this Laplacian approach with a traditional scalp reference, the results can, in principle, clarify the degree to which correlations with intelligence are driven by variation in activity of focal cortical sources (as revealed by the Laplacian) versus more aggregated activity that reflects contributions from deeper sources and greater signal mixing related to volume conduction.

In our study, we applied the surface Laplacian transform via the CSD toolbox (Kayser & Tenke, 2006; http://psychophysiology.cpmc.columbia.edu/Software/CSDtoolbox/) to the data and calculated amplitude and latency measurements afterwards. In the following (and in all Figures/Tables) we will refer to the former approach (described above) as ‘scalp-average referenced’ and to the latter as ‘Laplacian-transformed’.

In contrast to the scalp-averaged reference approach in which we used the same electrodes and time windows as Stefanics et al. (2011), for these analyses, we opted for a more data-driven approach that aligned with our sample. Thus, we first averaged the difference waves over all participants and conditions (*N* = 50 participants, six difference waves) to obtain a sample-specific grand-average difference wave. As depicted in Figure 2 (lower panels), we observed a clear negative maximum at electrodes POz and Oz between 125 and 275 ms post-stimulus, which were chosen as the electrode and time-windows for the individual difference analyses in the Laplacian-transformed data. As with the scalp-average referenced approach, the amplitude and latency values were extracted individually for each subject at POz and Oz separately, and averaged afterwards.

**Figure 2.**
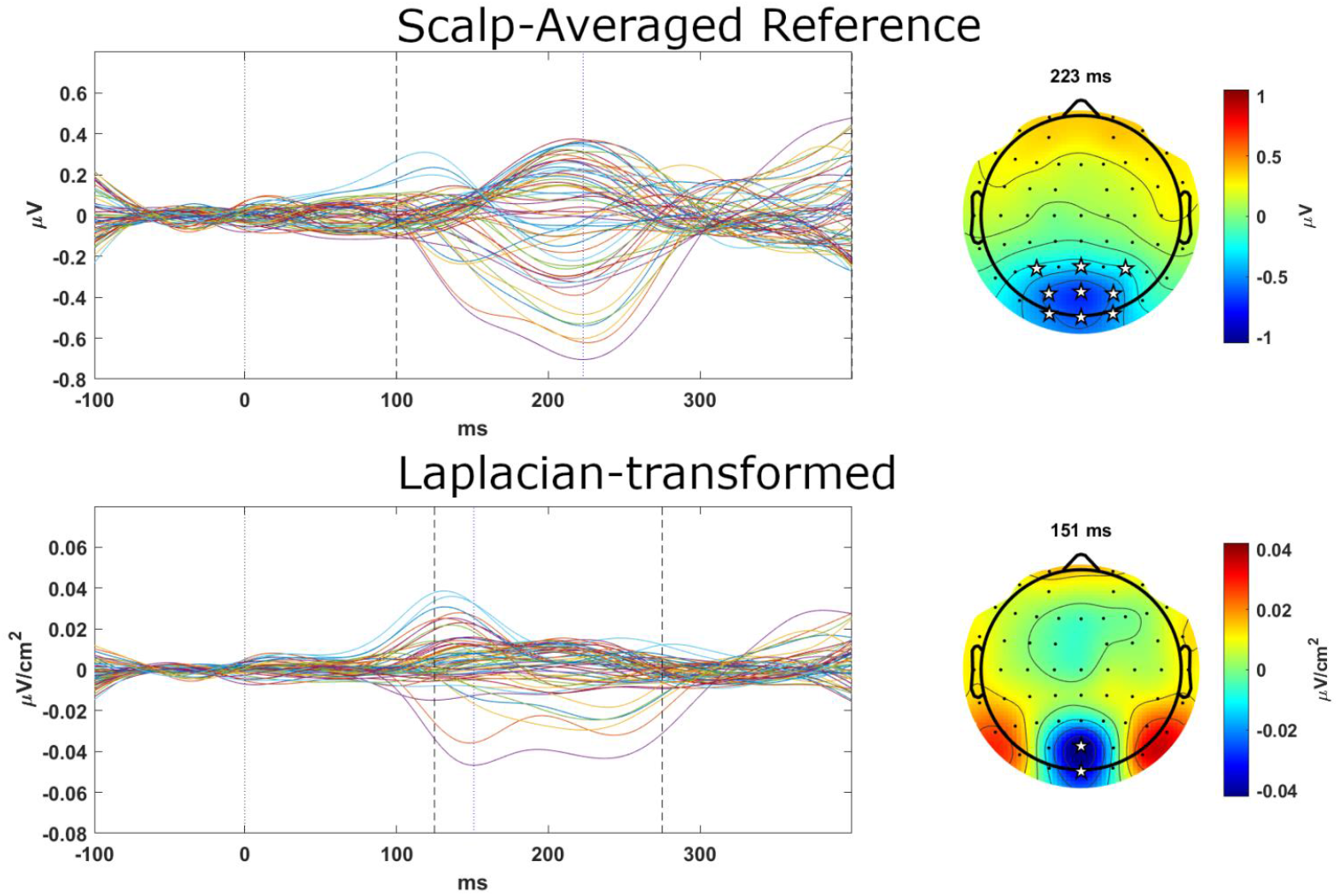
Grand-averaged ERP difference waves (MMNs) for scalp-averaged-referenced and Laplacian-transformed data. Top: Grand-averaged, scalp average-referenced difference waves and peak activity, over all participants and conditions. Bottom: Analogous plots for the Laplacian-transformed data. Left: Different channels are illustrated in different (randomly selected) colors. Dashed black lines indicate measurement windows for each measurement approach, blue dotted lines indicate peak ERP difference waves (vMMN) latencies depicted in the scalp maps at right. Right: Scalp maps. Stars highlight the electrodes that were used to calculate vMMN amplitudes and latencies for all individual difference analyses. Specifically, in the scalp-average referenced approach signed area amplitudes and 50% fractional area latency values were calculated in the same time window and on the same electrodes as in Stefanics et al. (2011), i.e., electrodes P3, Pz, P4, PO3, POz, PO4, O1, Oz, O2 and time window 100-400 post-stimulus. In the Laplacian-transformed approach, electrodes POz and Oz were used and the time window was set to 125-275 ms post-stimulus (identified in our sample; see Methods). In both approaches amplitude and latency values were calculated for each electrode separately and averaged across respective electrodes afterwards. Note that units and measurement scales differ: scalp-referenced data are in µV and Laplacian-transformed data are in µV/cm^2^.

Thus, the following analyses of condition effects involve only the scalp-average referenced data (pooling conditions across stimulus colors), whereas the analysis of associations between vMMN amplitudes/latencies and intelligence were conducted using the respective electrode-averaged values from the scalp-averaged reference and Laplacian-transformed difference waves. Grand-averaged difference waves, separated by condition and referencing scheme are presented in Figures 3 and 4.

**Figure 3.**
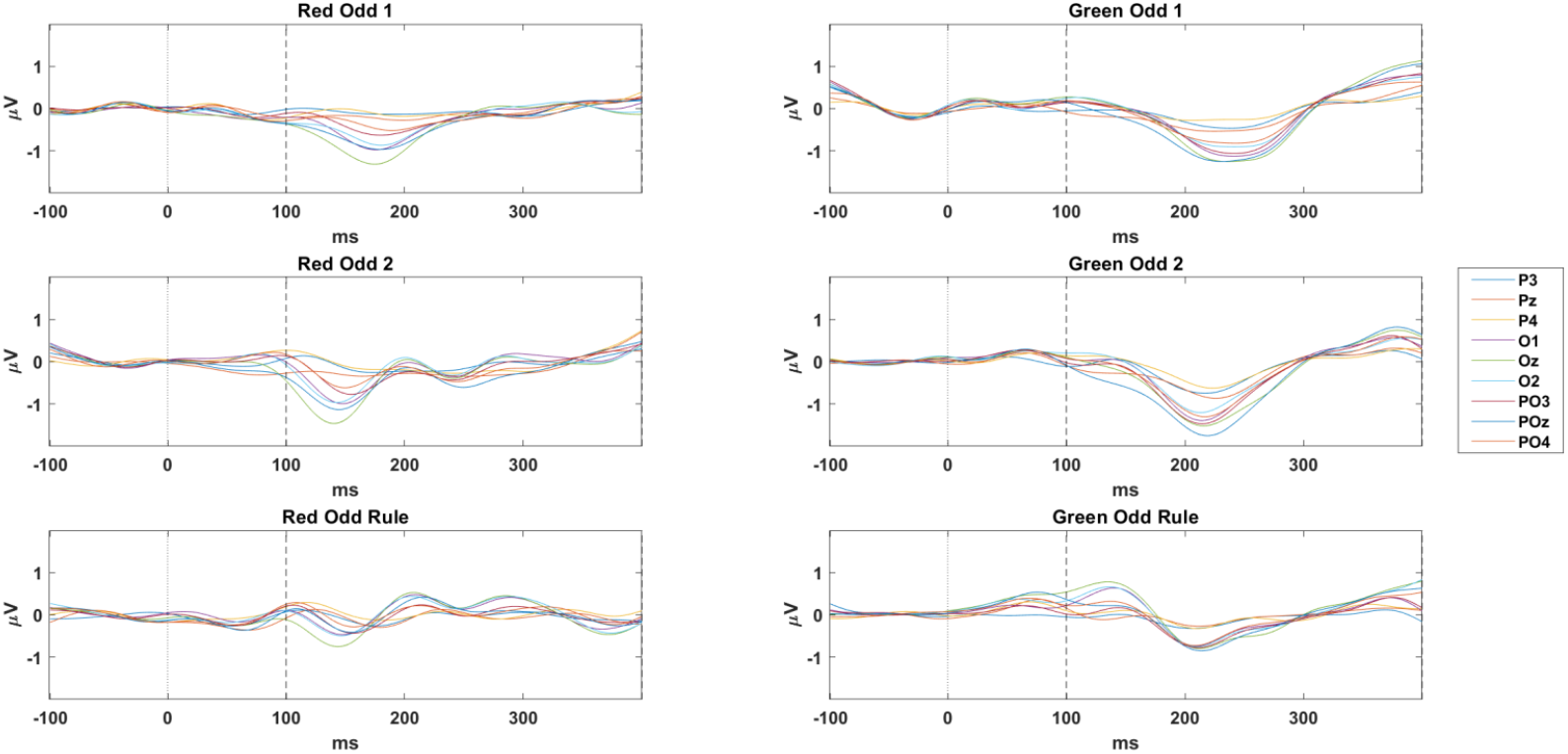
Plots depict the grand-averaged, scalp-averaged reference difference waves, over all participants, for each vMMN condition, in the nine electrodes of interest. Each difference wave was computed by subtracting the ERP elicited by standard stimuli from that elicited by deviant stimuli, in the same stimulus position and opposite color within the same task block. “Odd” designates the color of the deviant stimulus in that block. Dashed black lines indicate the measurement windows from 100-400 ms post-stimulus.

**Figure 4.**
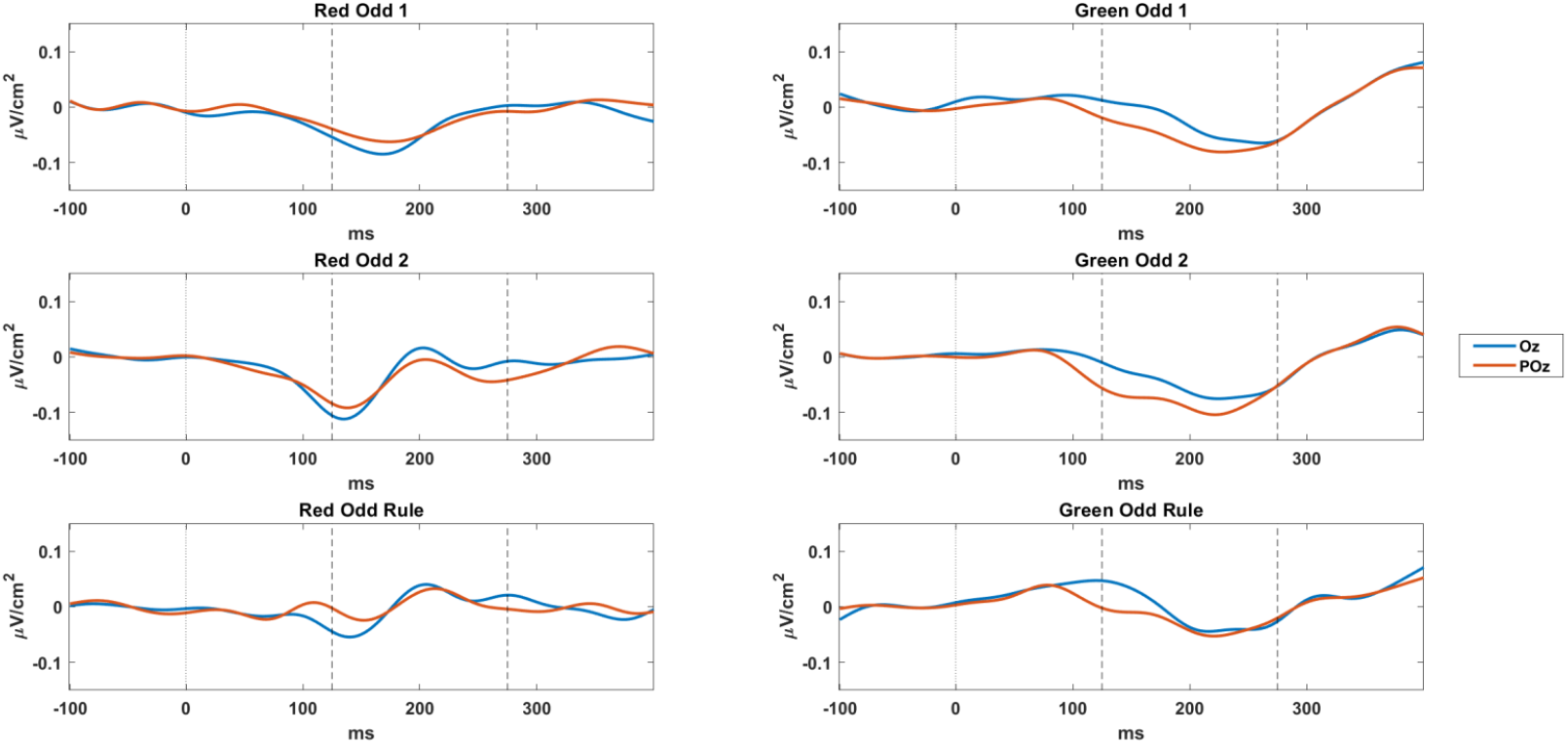
Plots depict the grand-averaged, Laplacian-transformed difference waves, over all participants, for each vMMN condition, in the two electrodes of interest. Each difference wave was computed by subtracting the ERP elicited by standard stimuli from that elicited by deviant stimuli, in the same stimulus position and opposite color within the same task block. “Odd” designates the color of the deviant stimulus in that block. Dashed black lines indicate the measurement windows from 125-275 ms post-stimulus.

#### Amplitude and Latency measures

For each subject-specific trial-averaged ERP and ERP difference wave (vMMN), we used ERPLAB algorithms to calculate signed integral area amplitudes and the 50% fractional area latencies (Luck, 2014). Signed area amplitude is obtained by taking the area of the shape that is produced between the boundary of the waveform (i.e., a positive or negative deflection) relative to the baseline voltage, within the time-window of interest (https://github.com/lucklab/erplab/wiki/ERP-Measurement-Tool). Since the MMN is defined as a relative negativity, the area for any positive deflections within the time-window were subtracted from the area for negative deflections. Unlike mean amplitude measures, area measures have the advantage of being less susceptible to noise and less affected by cross-study differences in time windows (Luck, 2014).

Fractional area latency represents an estimate of the midpoint latency, which is less sensitive to noise and thus more reliable than the frequently used ‘peak latency’ measure (Luck, 2014). The computation of the fractional area latency involves determining the area under the ERP waveform over a given time window and then finding the time point at which the area reaches a specified percentage of its peak (as recommended in Luck, 2014, we used 50%). Note that because the vMMN is defined as a relative negativity, we excluded instances in which no negative deflection was observed at all. Thus, the amplitude results reported below reflect the area of the negative deflections subtracted from any positive values, and similarly, fractional area latencies were calculated for negative deflections only.

### Data Analyses

Primary data analyses were conducted in SPSS version 25. We performed two sets of analyses with the goal of (1) verifying whether the experimental manipulation was successful in eliciting a vMMN in all conditions (i.e., simple and rule), and (2) assessing whether vMMN amplitudes or latencies correlate with individual differences in intelligence.

#### Effects of Stimulus Condition

For the within-subject analyses of condition effects we sought to replicate Stefanics et al. (2011) as closely as possible. Hence, following their approach, we collapsed over red and green stimuli and conducted two repeated-measures ANOVAs, with four factors (Stimulus Deviance: standard vs. deviant; Stimulus Position: first vs. second; Anteriority: parietal, parieto-occipital, occipital; and Hemisphere: left, midline, right; factorial design: 2 × 2 × 3 × 3) to examine the effects of experimental manipulations on ERP amplitudes and latencies. For the rule condition, all ERPs had been calculated on 2^nd^-position stimuli (see above), resulting in 2 × 3 × 3 ANOVAs (Deviance x Hemisphere x Anteriority) for both amplitudes and latencies. The Greenhouse-Geisser correction was applied for all tests containing three or more factors (Luck, 2014), and the Bonferroni correction was applied to control for multiple comparisons when resolving main effects.

Of note, we observed that in a small number of instances the trial-averaged ERP (within-subject analyses) or difference wave (individual difference analyses) was exclusively within the positive range in the respective time-window of interest. In these cases, the integral area amplitude takes a positive value, but the fractional area latency measure was not defined at all. Thus, the analyses of within-subjects effects (ANOVAs) included all *N* = 50 participants in respect to vMMN amplitudes, while within-subjects analyses conducted on latency measures relied on only *N* = 48 participants in the basic condition and on *N* = 47 participants in the rule condition after excluding those with positive difference waves.

#### Individual Difference Analysis

The individual difference analyses aimed to focus solely on the amplitudes and latencies of the six ERP difference waves. However, as we observed a reliable vMMN only in the four simple conditions but not in the rule condition (see Results), individual differences analyses were conducted for the four simple conditions only. To reduce the number of statistical comparisons, we decided *a priori* to average the amplitude/latency values computed from the relevant electrodes and time-windows in each referencing scheme. Thus, as described above, we averaged across amplitude and latency values in the nine electrodes (P3, Pz, P4, PO3, POz, PO4, O1, Oz) used in Stefanics et al. (2011), and across POz and Oz for the Laplacian-transformed approach. This produced one amplitude and one latency value for each participant for each of the two referencing schemes.

To address our primary question regarding the associations between vMMN amplitudes and latencies and intelligence, we then conducted partial correlations (corrected for age, sex, and participants’ minimum number of trials in any condition, as a proxy for ERP quality) between RAPM scores and amplitude/latency measures of each of the four ERP difference waves for each of the two measurement approaches. For the analyses of within subject effects of task condition, uncorrected *p*-values are reported for omnibus tests, whereas simple main effects have been adjusted via the Bonferroni correction. In contrast, for the individual difference analyses we report Bayes Factors instead of *p*-values to assess the significance of findings (see below).

A final methodological consideration concerns the individual difference analyses of the ERP latencies. As the calculation of the 50%-fractional area latency measure resulted in missing values in some cases (see above), the respective values could not simply be averaged across electrodes as it has been done for the amplitude measure (see Schafer & Graham, 2002). Thus, we conducted multiple imputation, which is a well-established principled method for dealing with missing data (Graham, 2009). In brief, the process involves replacing each missing value with a simulated value, which is obtained via a random draw from a predicted distribution of the respective variable (for details see Schafer & Graham, 2002). This process is performed multiple times, resulting in a set of complete datasets each containing simulated copies of the missing values. Of note, these datasets were generated in a way that the distributional properties of the variable were preserved. Each imputed dataset is then analyzed with conventional statistics (here partial correlations between RAPM score and latency value controlling for age, sex, and minimum number of trials in any condition), and the resulting parameter estimates (partial correlations, *r*) are pooled arithmetically over all imputed datasets (Sinharay et al, 2001). Following the guidelines reported in Sinharay et al. (2001), we generated ten imputed datasets. As the pooled partial correlation coefficients were aggregated through Fisher-Z-transform, we could not report conventional *p*-values in these cases.

To also quantify the evidence in favor of the null hypothesis (i.e., the absence of an association between intelligence and ERP difference waves measures) Bayes Factors (BF^01^, Jeffreys, 1998; Wagenmakers, 2007; Wetzels & Wagenmakers, 2012) and 95% credibility intervals (CIs) were calculated for all associations of primary interest. We used the Bayesian test for correlated pairs (Jeffreys, 1961) and the default prior (stretched beta distribution of width 1, also termed Jeffrey prior) as implemented in JASP version 0.11.1 (Love et al., 2015). To approximate pooled partial correlations (controlled for age, sex, and minimum trial numbers), we based these analyses on residual intelligence scores, i.e., any variance due to age, sex, and minimum trial numbers was regressed from RAPM sum scores prior to computation of Bayesian correlations.

### Data and Code Availability

All analysis code used in the current study has been deposited on GitHub at https://github.com/KirstenHilger/IQ_Coding. The raw study data can be accessed from the authors upon request.

## Results

Descriptive statistics for reaction times, accuracies, and RAPM performance are listed in Table 1. Participants were highly accurate in performing the target detection task (*M* = 97% correct; *SD* = 0.07). Descriptive statistics for ERP amplitudes and latencies are presented in Table 2 (for simplicity, values are presented only for electrode POz, averaged across red and green simple conditions). In terms of numerical effects (i.e., prior to conducting inferential statistics), examination of the relevant means in the simple conditions suggest that amplitudes were larger (more negative-going), and latencies were faster in response to second-position stimuli-relative to first-position stimuli, and in response to deviant stimuli relative to standard stimuli. No clear differences were evident for amplitudes or latencies in the rule condition. Raw ERPs for the 12 experimental sub-conditions are presented in Supplementary Figures S1 and S2 for the scalp-averaged reference and Laplacian transformed data, respectively.

**Table 2.**
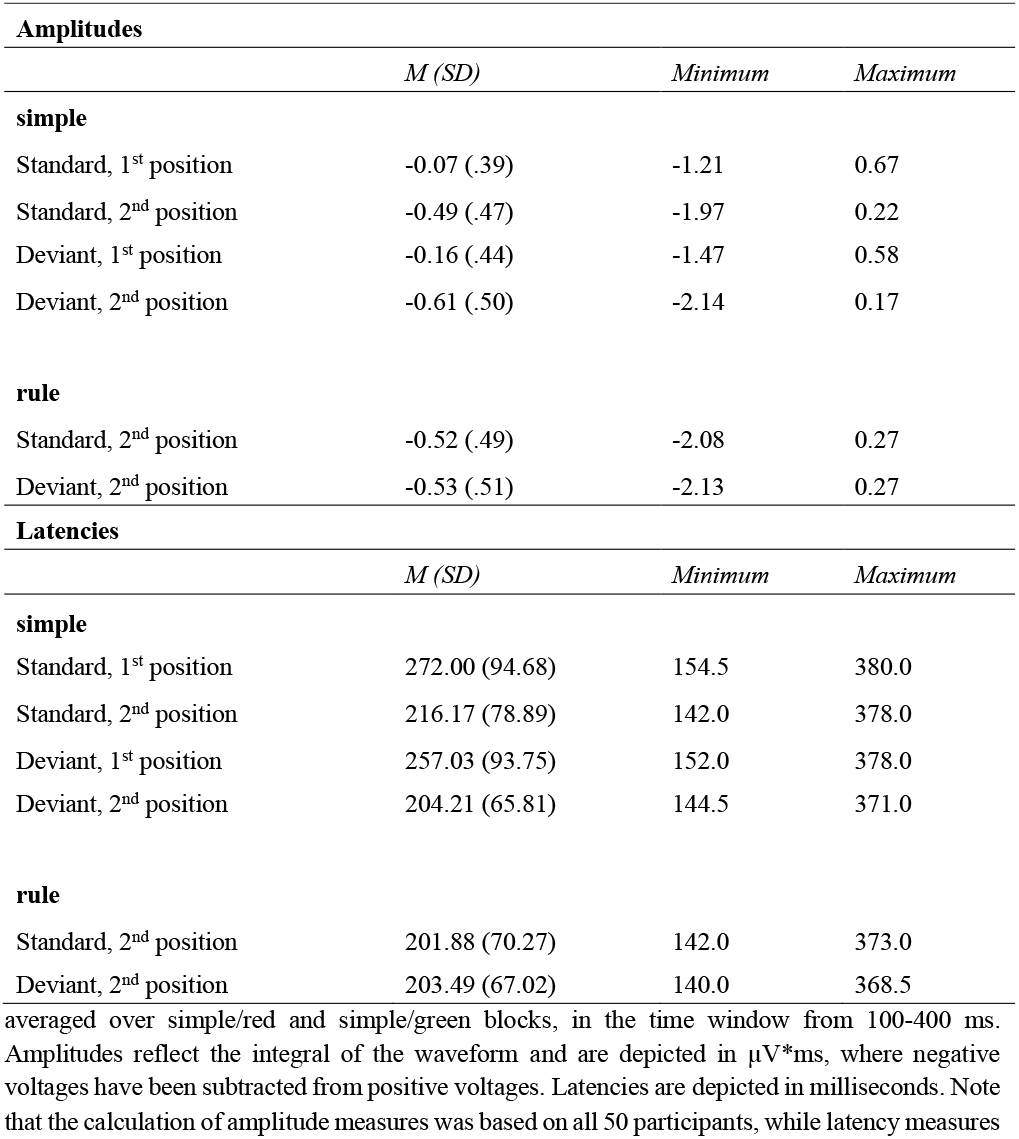
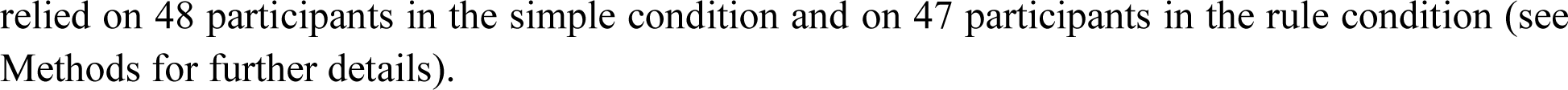
Descriptive statistics for ERP amplitudes and latencies based on the scalp-average referenced approach

| **Amplitudes** |  |  |  |
| --- | --- | --- | --- |
|  | *M (SD)* | *Minimum* | *Maximum* |
| **simple** |  |  |  |
| Standard, 1<sup>st</sup> position | -0.07 (.39) | -1.21 | 0.67 |
| Standard, 2<sup>nd</sup> position | -0.49 (.47) | -1.97 | 0.22 |
| Deviant, 1<sup>st</sup> position | -0.16 (.44) | -1.47 | 0.58 |
| Deviant, 2<sup>nd</sup> position | -0.61 (.50) | -2.14 | 0.17 |
| **rule** |  |  |  |
| Standard, 2<sup>nd</sup> position | -0.52 (.49) | -2.08 | 0.27 |
| Deviant, 2<sup>nd</sup> position | -0.53 (.51) | -2.13 | 0.27 |
| **Latencies** |  |  |  |
|  | *M (SD)* | *Minimum* | *Maximum* |
| **simple** |  |  |  |
| Standard, 1<sup>st</sup> position | 272.00 (94.68) | 154.5 | 380.0 |
| Standard, 2<sup>nd</sup> position | 216.17 (78.89) | 142.0 | 378.0 |
| Deviant, 1<sup>st</sup> position | 257.03 (93.75) | 152.0 | 378.0 |
| Deviant, 2<sup>nd</sup> position | 204.21 (65.81) | 144.5 | 371.0 |
| **rule** |  |  |  |
| Standard, 2<sup>nd</sup> position | 201.88 (70.27) | 142.0 | 373.0 |
| Deviant, 2<sup>nd</sup> position | 203.49 (67.02) | 140.0 | 368.5 |

averaged over simple/red and simple/green blocks, in the time window from 100-400 ms. Amplitudes reflect the integral of the waveform and are depicted in $\mu\text{V}\cdot\text{ms}$ , where negative voltages have been subtracted from positive voltages. Latencies are depicted in milliseconds. Note that the calculation of amplitude measures was based on all 50 participants, while latency measures

### Effects of Stimulus Condition

#### ERP Amplitudes – Simple Conditions

The four-way ANOVA examining the effects of stimulus Position (1^st^ or 2^nd^), Deviance (standard vs. deviant), Anteriority (parietal, parieto-occipital, occipital), and Hemisphere (left, midline, right) revealed significant main effects of Position (*F*(1, 49) = 157.908, *p* <. 001, partial *η*^2^ = .763), Deviance (*F*(1, 49) = 50.252, *p* <. 001, partial *η*^2^ = .506), and Hemisphere (*F*(1.548, 75.846) = 38.898, *p* <. 001, partial *η*^2^ = .443; see also Supplementary Table S1).

More detailed simple main effect tests revealed first that ERP amplitudes were generally more negative for second-relative to first-position stimuli (see Figure S3), with more negative ERPs at midline relative to left or right hemisphere sites (all *p*-values < .001), which did not differ from each other (all *p*-values ≥ .164). In addition, amplitudes were larger at parieto-occipital sites than either parietal or occipital sites in the left hemisphere (all *p*-values ≤ .014), and at the midline, (all *p*-values ≤ .003), with a similar pattern in the right hemisphere (all *p*-values ≤ .060). There were no differences between left and right hemisphere sites at any level of Anteriority (all *p*-values ≥ .156).

Second, we observed that ERP amplitudes were consistently larger (more negative) at midline relative to left or right hemisphere sites across parietal, parieto-occipital, and occipital electrodes (all *p*-values < .001), whereas left and right hemisphere sites did not differ from each other at any level of Anteriority (all *p*-values ≥ .820; and see Supplementary Figure S3). The only mean difference to emerge in the anterior-posterior dimension was a larger effect at POz relative to Oz (*p* = .007; all other *p*-values ≥ .231).

Third, the interactions of Deviance and Hemisphere and Deviance and Anteriority (see Supplementary Table S1) indicate that, consistent with the key assumption of MMN research, deviant stimuli elicited larger ERP amplitudes than standards at all levels of Hemisphere (all *p*-values > .001) and at all levels of Anteriority (all *p*-values > .001). When the effects of stimulus Deviance were considered with respect to their spatial dimensions, they also suggest a maximum near POz (Supplementary Figure S4). Specifically, amplitudes were consistently larger at the midline than in the left or right hemisphere for both deviant and standard stimuli (all *p*-values < .001), while amplitudes over left and right hemispheres did not differ from each other (all *p*-values ≥ .639); and likewise, deviant stimuli elicited greater amplitudes at parieto-occipital than either parietal (*p* = .041) or occipital sites (*p* = .016), which did not themselves differ from each other (*p* > .999). This difference was not observed for standard stimuli, however, where amplitudes were larger at POz than Oz (*p* = .010), but not Pz (*p* = .327), and similarly, Pz and Oz did not differ from one another (*p* = .999). For the sake of interest, Supplementary Table S1 contains a full statistical analysis of all possible interactions.

#### ERP Amplitudes – Rule Condition

In the rule condition, we examined the effects of stimulus Deviance, Hemisphere, and Anteriority, and their respective interactions. Only the main effects of Anteriority (*F*(1.285, 62.985) = 7.608, *p* = .004, partial *η*^2 =^ .134), Hemisphere (*F*(1.576, 77.217) = 38.754, *p* < .001, partial *η*^2 =^ .442), and their interaction reached statistical significance (*F*(2.997, 146.872) = 6.861, *p* < .001, partial *η*^2 =^ .123). Thus, the experimental manipulation in the rule condition did not elicit a clear vMMN, i.e., there was no significant difference in group-averaged amplitudes between deviant and standard stimuli (*F*(1,49) = .232, *p* = .632, partial *η*^2 =^ .005). As with the results for the pooled simple stimuli (see above), the significant interaction reflected the spatial specificity of the ERPs, such that amplitudes were largest near POz (moving both right to left and anterior to posterior), with particularly pronounced differences between POz (midline) and PO3/PO4 (left/right) and relative to analogous contrasts at parietal or occipital sites. Supplementary Table S2 lists the full range of statistics.

In summary, ERP amplitude results for the simple conditions were consistent with previous research indicating larger ERPs in response to deviant versus standard stimuli and following the second versus the first stimuli of a pair. Although stimulus Position and Deviance did not interact, they showed similar spatial effects, with each being most pronounced near the midline parieto-occipital electrode POz. In contrast to a previous study using the same paradigm (Stefanics et al., 2011), in our sample, the rule condition deviant stimuli did not elicit larger amplitudes than standard stimuli.

#### ERP Latencies – Simple Conditions

In the four-way ANOVA examining the effects of Stimulus Position, Deviance, Anteriority, and Hemisphere on ERP latencies, the main effects of Position (*F*(1, 47) = 67.577, *p* < .001, partial *η*^2^ = .590), Deviance (*F*(1, 47) = 16.443, *p* < .001, partial *η*^2^ = .259), and Hemisphere (*F*(1.668, 78.375) = 12.693, *p* < .001, partial *η*^2^ = .213) reached statistical significance, as well as the two-way interactions of Position and Deviance (*F*(1,47) = 6.515, *p* = .014, partial *η*^2^ = .122), Position and Hemisphere (*F*(1.903, 89.432) = 5.429, *p* = .007, partial *η*^2^ = .104), and Anteriority by Hemisphere (*F*(2.749, 129.195) = 3.853, *p* = .013, partial *η*^2^ = .076). All three- and four-way interactions were not significant (see also Supplementary Table S3).

More detailed simple main effect tests on the latencies indicated first that ERP latencies were consistently earlier for second-vs. first-position stimuli (all *p* < .001). This pattern was found across all two-way interactions such that the various interactions moderated this effect. Second, as indicated by the interaction of Position by Hemisphere, latencies were consistently earlier at midline relative to left or right hemisphere electrodes (all *p* ≤ .05), which did not differ from each other (all *p* > .999); though this effect was greater for first relative to second-position stimuli. Last, in respect to the interaction between Hemisphere and Anteriority, we observed that latencies were shorter at midline relative to right and left hemisphere sites (all *p* ≤. 013), that right and left sites did not differ from each other (all *p* >. 999), and that this difference was especially pronounced between midline and right parietal and parieto-occipital sites (see Supplementary Figure S5). A full list of statistics is presented in Supplementary Table S3. As no clear vMMN was observed in the rule condition (see above), latencies were not analyzed here.

In summary, analyses of ERP latencies from the simple conditions indicated that, consistent with previous research, latencies tended to be faster to deviant than to standard stimuli, to second-relative to first position stimuli, and to midline relative to peripheral electrodes. The latter became most visible at parieto-occipital sites i.e., especially at POz (see Supplementary Figure S5).

### Individual Differences Analyses

#### Visual Mismatch Negativity and Individual Differences in Intelligence

In respect to our main question of interest, i.e., whether intelligence is associated with neural indicators of simple visual discrimination processes (i.e., the vMMN), we observed only negligible to small correlations for the amplitude measures (interpretation in accordance with Cohen, 1988), which varied between .01 (Laplacian transformed: Green Odd 1) and -.14 (scalp-average referenced: Red Odd 1; Table 3). All Bayes Factors (BF_01_) were above three, providing substantial evidence for the null hypothesis, i.e., absence of association between intelligence and MMN amplitudes (interpretation of BFs in accordance with Jeffreys, 1961 and Wagenmakers et al., 2011). As an example, the BF_01_ of 5.54 for the association between intelligence and the vMMN amplitude for Green first-position Odd difference wave (Table 3) indicates that the null hypothesis is ∼ six times more likely than the alternative hypothesis (i.e., presence of a correlation). Also, six out of eight 95% credibility intervals included zero, which further supports the absence of an association in most cases. For the sake of completeness, partial correlations between vMMN amplitudes of different conditions are provided in Supplementary Table S4.

**Table 3.** Partial correlations, Bayes Factors, and 95% credibility intervals for associations between intelligence and vMMN amplitudes for both measurement approaches

| **Scalp-Averaged Reference** |  |  |  |  |
| --- | --- | --- | --- | --- |
| Statistic | Green Odd 1 | Green Odd 2 | Red Odd 1 | Red Odd 2 |
| *Partial r* | .03 | .13 | -.14 | .02 |
| *BF<sub>01</sub>* | 5.54 | 3.92 | 3.85 | 5.65 |
| *CI* | [-.05, .12] | [.04, .21] | [-.21, -.05] | [-.07, .10] |
| **Laplacian-Transformed** |  |  |  |  |
| *Partial r* | .01 | .07 | -.01 | -.03 |
| *BF<sub>01</sub>* | 5.66 | 5.12 | 5.65 | 5.54 |
| *CI* | [-.26, .28] | [-.21, .33] | [-.28, .26] | [-.30, .24] |

*Note:* In the scalp-average referenced approach signed area amplitudes were calculated in the same time window and on the same electrodes as in Stefanics et al. (2011), i.e., electrodes P3, Pz, P4, PO3, POz, PO4, O1, Oz, O2 and time window 100-400 post-stimulus. In the Laplacian-transformed approach, electrodes POz and Oz were used and the time window was set to 125-275 ms post-stimulus (identified in our sample; see Methods). In both approaches signed area amplitudes were calculated for each electrode separately and averaged across respective electrodes afterwards. Partial $r$ = pooled partial correlation coefficients controlling for variance due to age, sex, and minimum number of artifact-free trials. Green Odd 1 = ERP difference wave calculated from red standards minus green deviants in the first position of the stimulus pairs in the simple/green block, etc. “Odd” always refers to the color of the deviant stimulus in the relevant condition/stimulus position. Bayes Factors ( $BF_{01}$ ) in favor of the null hypothesis (i.e., absence of correlation) and 95% credibility Intervals (CI) are depicted for associations with RAPM scores.

With respect to the latencies for the scalp-averaged reference data, pooled correlations between intelligence and vMMN were again of negligible to small effect sizes (Cohen, 1988) and varied between -.02 (Green Odd 1 and Red Odd 2) and .12 (Green Odd 2; see Table 4). All Bayes Factors were larger than three indicating substantial evidence for the null hypothesis and three out of four credibility intervals include zero. For the Laplacian-transformed data, pooled correlations were also of negligible (Green Odd 2, *r* = -.05) to small (e.g., Green Odd 1, *r* = .24) effect sizes. However, only the Bayes Factors for Green Odd 2 and Red Odd 2 were greater than three, providing substantial evidence for the null hypothesis. In contrast, the Bayes Factors for Green Odd 1 and Red Odd 1 fell between one and three providing anecdotal evidence for the null hypothesis and suggest a need for further investigation. Only the credibility interval of Green Odd 2 included zero, while all other were slightly (Red Odd 2) or clearly (Green Odd 1, Red Odd 1) distinct from zero (Table 4). Partial correlations between vMMN latencies of different conditions are provided in Supplementary Table S5. Scatterplots depicting the association between vMMN amplitude and latency variables and RAPM scores are presented in Supplementary Figures 6 and 7.

**Table 4.** Pooled partial correlations, Bayes Factors, and 95% credibility intervals for associations between intelligence and vMMN latencies for both measurement approaches

| **Scalp-Averaged Reference** |  |  |  |  |
| --- | --- | --- | --- | --- |
| Statistic | Green Odd 1 | Green Odd 2 | Red Odd 1 | Red Odd 2 |
| *Partial r* | -.02 | .12 | .06 | -.02 |
| *BF<sub>01</sub>* | 5.51 | 4.02 | 5.10 | 5.57 |
| *CI* | [-.10, .07] | [.03, .20] | [-.03, .14] | [-.10, .07] |
| **Laplacian-Transformed** |  |  |  |  |
| *Partial r* | .24 | -.05 | .20 | .12 |
| *BF<sub>01</sub>* | 1.46 | 5.33 | 2.37 | 4.27 |
| *CI* | [.16, .32] | [-.13, .04] | [.11, .28] | [.03, .19] |

*Note:* In the scalp-average referenced approach 50% fractional area latency values were calculated in the same time window and on the same electrodes as in Stefanics et al. (2011), i.e., electrodes P3, Pz, P4, PO3, POz, PO4, O1, Oz, O2 and time window 100-400 post-stimulus. In the Laplacian-transformed approach, electrodes POz and Oz were used, and the time window was set to 125-275 ms post-stimulus (identified in our sample; see Methods). In both approaches 50% fractional area latency values were calculated for each electrode separately and averaged across respective electrodes afterwards. Partial $r$ = pooled partial correlation coefficients controlling for variance due to age, sex, and minimum number of artifact-free trials. Green Odd 1 = ERP difference wave calculated from red standards minus green deviants in the first position of the stimulus pairs in the simple/green block, etc. “Odd” always refers to the color of the deviant stimulus in the relevant condition/stimulus position. Bayes Factors ( $BF_{01}$ ) in favor of the null hypothesis (i.e., absence of correlation) and 95% credibility Intervals (CI) are depicted for associations with RAPM scores.

## Discussion

This study presents a systematic test for a relation between intelligence and the MMN as an index of basic, pre-attentive automatic discrimination processes in the visual domain. Using a paradigm that has previously been shown to elicit a vMMN (Stefanics et al., 2011), we first replicated within-subject effects elicited by the experimental manipulation. Then, we tested for associations between individual differences in intelligence and vMMN amplitudes and latencies. However, only negligible to small correlations were observed. Bayes Factors suggest substantial evidence for the absence of associations between vMMN variables and intelligence in most cases, and we observed critical dependency of the results on different measurement approaches.

The within-subject effects of stimulus position, perceptual deviance, and scalp location, in addition to violations of more abstract, rule-based expectations were examined first. For the more basic perceptual manipulation (simple condition), our results replicate previous findings for ERP amplitudes. Specifically, we observed a clear vMMN that was maximal near parietal-occipital, midline electrodes (Figure. 2), and larger (more negative) amplitudes in response to deviant and 2^nd^-position stimuli, relative to standard stimuli or those presented 1^st^ in a pair. Although the rule condition also elicited an ERP with the expected parieto-occipital topography, there were no amplitude effects related to stimulus deviance. Analyses of vMMN latencies in the simple condition revealed that neural activity peaked more rapidly in response to 2^nd^ versus 1^st^-position stimuli and was also faster for deviants relative to standards. Also here, the spatial effects were centered near POz.

In the individual differences analyses probing for relations between intelligence and vMMN amplitudes or latencies, we compared two different referencing schemes—the scalp-averaged reference and current source density via the Laplacian transform. For the scalp-averaged reference data, we relied on an *a priori* measurement window and set of electrodes, whereas, for the Laplacian-transformed data, measurement parameters were optimized to the current sample in a data-driven approach. As expected, the Laplacian transform revealed a more focal pattern of neural activity, which also peaked earlier than in the scalp-referenced data (based on the grand-average). Notwithstanding, both analyses approaches revealed only negligible to small relationships between vMMN amplitudes and performance on the RAPM, and Bayes Factors suggested substantial evidence for the absence of any association. Correlations between vMMN latencies and intelligence likewise exhibited only negligible to small effect sizes. However, the corresponding Bayes Factors for these associations showed important differences across the two measurement approaches. Specifically, for the Laplacian-transformed data, there was only weak evidence for the null hypothesis in some conditions. For first position deviant stimuli, in both the Red and Green blocks, we observed small positive correlations, with 95% credibility intervals exclusively in the positive range. Thus, these data suggest a small positive relationship between intelligence and some vMMN latencies under the current approach.

On the one hand, this could suggest that the improved spatial resolution provided by the Laplacian-transform, and the more sample-specific approach to determine measurement parameters, were somewhat more successful in capturing the neural sources most relevant to the paradigm and perhaps to individual differences. Some additional support for this assumption comes from the fact that, although the sign of these effects contradicts the general finding of *negative* associations between ERP latencies and intelligence, there is nevertheless some evidence for positive associations between intelligence and the latencies of early ERPs (Schubert, 2017). On the other hand, insofar as the individual differences effects were generally small and inconsistent with the larger literature, they require replication and merit only a tentative interpretation.

The current findings contrast with studies reporting correlations between intelligence and the MMN in the auditory domain. Although results for individual studies are mixed, the overarching picture suggests larger MMN amplitudes (Sculthorpe et al., 2009; Troche et al., 2009, 2010), faster MMN latencies (e.g., Bazana & Stelmack, 2002; Beauchamp & Stelmack, 2006), or both (De Pascalis & Varriale, 2012; De Pascalis et al., 2014; Houlihan & Stelmack, 2012), in higher ability participants. At least one other study also examined the relation between intelligence and a version of the vMMN. That study, however, focused on socio-emotional rather than on automatic perceptual processes, and investigated amplitudes only. Specifically, the authors examined the vMMN in response to perceptually complex stimuli of neutral versus happy or sad facial expressions and observed larger MMN amplitudes in higher ability participants (Liu et al., 2015).

Plausible explanations for the discrepancy between the current results and previous studies include differences in measurement approaches, differences in specific task features, and broader considerations pertaining to different demands between the visual and auditory modalities. For example, although the current paradigm attempts to control participants’ attention via the manipulation of fixation cross length, the effectiveness of this manipulation is somewhat unclear. In visual MMN paradigms, both the attended and unattended stimuli are presented in the same modality. In contrast, in auditory MMN paradigms, participants’ attention is most often kept by tasks in a different domain such as vision (e.g., by reading or watching a silent video). As vision is humans’ dominant modality, efforts to control attention might be more effective in this latter case. A related consideration concerns potential differences in the degree to which the two modalities may admit of individual differences in early perceptual processing. For example, it has been argued that auditory signals present an especially complex information processing challenge, due to their inherent transience, and the frequent need to distinguish multiple objects from within a single, multi-layered input (Fitzgerald & Todd, 2020). Insofar as the two sensory modalities evolved to cope with different features of the environment, it is interesting to consider whether anatomical or physiological differences between early auditory and early visual pathways may affect the degree to which each admits of individual differences in their response properties. If so, this may in turn affect their potential for relations with intelligence. Similar considerations may also help to explain the discrepancy between our study that manipulated the colors and presented order of arrays of circles, versus that of Liu et al. (2015), which elicited the vMMN in response to emotional faces, i.e., to rather complex and highly socially salient stimuli.

A final consideration relates to the ‘expected’ effect sizes for relations with intelligence in vMMN and similar paradigms, especially in light of the general null findings. The first possibility is that there may simply not exist a relation between intelligence and early visual processing, for the reason that intelligence is fundamentally concerned with higher-order cognitive and neural capacities. An alternative view is that while intelligence is best *indexed* by higher-order processes, as a hierarchical construct (i.e., that captures aggregated variability across many functions), its relations to cognitive and neural processes may systematically vary as a function of task parameters like complexity or uncertainty (reviewed in Euler, 2018). If so, it should perhaps be expected that relations between neural variables and intelligence will be small in ‘basic’ perceptual paradigms. These relations may increase as greater task demands recruit broader neural networks, which may admit of greater individual differences (e.g., see Mueller et al., 2012). As such, future studies in this area would likely benefit from implementing a broader range of different task complexities (Euler & Schubert, 2021).

The present study also has several limitations which need to be considered. First, our sample size was comparatively small for analyses of individual differences in general (Yarkoni & Braver, 2010; Gignac, & Szodorai, 2016). Although the sample size was sufficient to detect medium to large effects (i.e., correlations > .35, see Methods) with 80% statistical power, we of course cannot exclude the possibility that, per the above, relations between intelligence and vMMN factors were smaller than anticipated, making this study under-powered to detect them. Nevertheless, the present data—which tested associations for both amplitudes and latencies, across four ERP difference waves and two referencing schemes—suggests that any relationship, if it exists, is apt to be rather small (and perhaps smaller than would be expected based on the previous literature on intelligence and auditory MMN). Replication studies using larger sample sizes are therefore essentially required. A second limitation concerns the rather low mean IQ in our sample, which we assume to be the result of using a relatively difficult test under a strict time limitation of 40 minutes, which hindered many participants’ ability to finish and, thus, increased heterogeneity. Third, in some deviant conditions, some individual participants had relatively few trials, potentially affecting the resulting ERPs. Although, it has been shown that reliable ERPs can be derived from fairly small numbers of trials (e.g., 6-8 trials in Olvet & Hajcak, 2009; 8 trials in Boudewyn et al., 2017; < 14 trials in Larson et al., 2010), it might be a general problem in MMN research that deviant and standard ERPs include different numbers of trials and might thus be characterized by different signal-to-noise ratios (SNR). For individual difference research, this issue is only problematic when differences in deviant vs. standard SNRs correlate systematically with the individual difference measure of interest. Nevertheless, we controlled our analyses for individual differences in the number of artefact-free trials. It may be worthwhile to develop methodological frameworks and experimental paradigms that equal the numbers of trials and to prevent this potential confound. Fourth, the choice to examine correlations of intelligence with ERP difference waves, rather than their constituent ERPs, may represent another limitation. Several recent papers have re-focused attention on psychometric problems associated with using difference scores in individual differences research (e.g., Hedge et al., 2018), including in ERP studies *per se* (Meyer et al., 2020). Although this appears not to have hindered most auditory MMN studies, given the variable and often low correlations between the difference waves themselves in the current data (Supplementary Tables S4-S5), this is a worthwhile possibility to investigate in future research. Finally, it has also been recently argued that EEG and ERP studies may be more apt to observe relations with intelligence when all variables are modelled as latent factors, and at comparable levels of psychological and physiological aggregation (Euler & Schubert, 2021; Schubert et al., 2017, 2021). While limitations of the present data preclude a more complex modelling approach along these lines, this represents another promising direction for future work.

In summary, this study presents a systematic test for a relationship between intelligence and a basic perceptual manipulation of the MMN in the visual modality. Using a previously validated paradigm, our results replicated prior experimental effects concerning the sensitivity of neural responses to deviant and second-position stimuli, albeit not to the violation of a more abstract, rule-based stimulus pattern. Most importantly, no associations between intelligence and vMMN amplitudes were observed and Bayesian analyses indicated substantial evidence against any relationship. In contrast, the associations between intelligence and vMMN latencies vary critically between different measurement approaches strongly requiring further investigation. Future studies in this area will benefit from larger sample sizes, from systematic comparisons of different measurement approaches and processing parameters, and from systematically exploring effects of task complexity and modality on relations between the MMN and intelligence.

## Acknowledgement

The authors thank Prof. Dr. Christian Fiebach, Prof. Dr. Ulrike Basten, and Dr. Jona Sassenhagen for support in acquiring ethics approval, data acquisition and for helpful comments to a previous version of the manuscript.

## Notes

https://github.com/KirstenHilger/IQ_Coding

### Competing Interest Statement

The authors have declared no competing interest.

### Summary of Updates

Beyond more detailed edits to improve the quality of our paper, we changed: - A more comprehensive elaboration on the motivation of our study and a stringer focus on the question how, in our opinion, research on the visual mismatch negativity (vMMN) can enhance our understanding about intelligence. - We re-run all of our analyses with subtracting the deviant from the standard of the very same trial and with the numbers of artefact-free EEG trials as additional control variable. - Figures and Tables were updated accordingly and we added new Figures both in the manuscript and the Supplement to enhance the clarity of our results.

