## Supplement for "Intelligence and Visual Mismatch Negativity: Is Pre-Attentive Visual Discrimination Related to General Cognitive Ability?"

**Supplementary Table S1**

Within-subjects ANOVA of simple conditions group-averaged ERP amplitudes by stimulus factors and electrode location

| Variable | *df* | *F* | *partial* $\eta^2$ | *p* |
|---|---|---|---|---|
| Position | 1, 49 | 157.91 | .76 | < .001** |
| Deviance | 1, 49 | 50.25 | .51 | < .001** |
| Anteriority | 1.30, 63.52 | 2.62 | .05 | .102 |
| Hemisphere | 1.55, 75.85 | 38.90 | .44 | < .001** |
| Position * Deviance | 1, 49 | 0.47 | .01 | .495 |
| Position * Anteriority | 1.27, 62.01 | 55.58 | .53 | < .001** |
| Position * Hemisphere | 1.66, 81.37 | 8.76 | .15 | .001* |
| Deviance * Anteriority | 1.68, 82.27 | 28.40 | .37 | <.001** |
| Deviance * Hemisphere | 1.22, 59.85 | 11.75 | .19 | .001* |
| Anteriority * Hemisphere | 3.12, 152.62 | 4.82 | .09 | .003* |
| Position * Deviance * Anteriority | 1.60, 78.26 | 1.85 | .04 | .171 |
| Position * Deviance * Hemisphere | 1.37, 67.19 | 1.05 | .02 | .333 |
| Position * Anteriority * Hemisphere | 2.96, 145.21 | 7.75 | .14 | <.001** |
| Deviance * Anteriority * Hemisphere | 3.08, 150.74 | 2.26 | .04 | .082 |
| Position * Deviance * Anteriority * Hemisphere | 3.28, 161.05 | 0.03 | <.01 | .995 |

*Note:* $N = 50$. Degrees of freedom are reported using the Greenhouse-Geisser correction for all tests containing three or more factors. Significant effects are marked with an asterisk (* = $p < .05$, ** = $p < .001$).

**Supplementary Table S2**

Within-subjects ANOVA of rule condition group-averaged ERP amplitudes by stimulus factors and electrode location

| Variable | *df* | *F* | *partial* $\eta^2$ | *p* |
|---|---|---|---|---|
| Deviance | 1, 49 | 0.23 | <.01 | .632 |
| Anteriority | 1.29, 62.99 | 7.61 | .13 | .004* |
| Hemisphere | 1.58, 77.22 | 38.76 | .44 | < .001** |
| Deviance * Anteriority | 1.45, 71.08 | 1.17 | .02 | .303 |
| Deviance * Hemisphere | 1.31, 64.06 | 2.00 | .04 | .158 |
| Anteriority * Hemisphere | 3.00, 146.87 | 6.86 | .12 | <.001** |
| Deviance * Anteriority * Hemisphere | 3.05, 149.48 | 0.56 | .01 | .644 |

*Note:* $N = 50$. Degrees of freedom are reported using the Greenhouse-Geisser correction for all tests containing three or more factors. Significant effects are marked with an asterisk (* = $p < .05$, ** = $p < .001$).

**Supplementary Table S3**

Within-subjects ANOVA of simple conditions group-averaged latencies by stimulus factors and electrode location

| Variable | *df* | *F* | *partial* $\eta^2$ | *p* |
|---|---|---|---|---|
| Position | 1, 47 | 67.58 | .59 | < .001** |
| Deviance | 1, 47 | 16.44 | .26 | < .001** |
| Anteriority | 1.29, 60.61 | 1.78 | .04 | .186 |
| Hemisphere | 1.67, 78.38 | 12.69 | .21 | < .001** |
| Position * Deviance | 1, 47 | 6.52 | .12 | .014* |
| Position * Anteriority | 1.55, 72.89 | 2.33 | .05 | .117 |
| Position * Hemisphere | 1.90, 89.43 | 5.43 | .10 | .007* |
| Deviance * Hemisphere | 1.96, 95.26 | 1.84 | .04 | .166 |
| Deviance * Anteriority | 1.45, 68.11 | 0.27 | .01 | .693 |
| Anteriority * Hemisphere | 2.75, 129.20 | 3.85 | .08 | .013* |
| Position * Deviance * Anteriority | 1.75, 82.11 | 0.93 | .02 | .387 |
| Position * Deviance * Hemisphere | 1.94, 90.92 | 3.05 | .06 | .054 |
| Position * Anteriority * Hemisphere | 2.92, 137.39 | 1.94 | .04 | .128 |
| Deviance * Anteriority * Hemisphere | 3.50, 164.47 | 0.78 | .02 | .523 |
| Position * Deviance * Anteriority * Hemisphere | 3.01, 141.31 | 0.69 | .01 | .562 |

*Note:* $N = 48$. Degrees of freedom are reported using the Greenhouse-Geisser correction for all tests containing three or more factors. Significant effects are marked with an asterisk (* = $p < .05$, ** = $p < .001$).

**Supplementary Table S4**

Pooled partial correlations for ERP difference waves (vMMN) amplitudes

| **Scalp Averaged Referenced** | | | | | |
|---|---|---|---|---|---|
| Variable | Green Odd 2 | Red Odd 1 | Red Odd 2 | Red Odd Rule | Green Odd Rule |
| Green Odd 1 | .37* | -.20 | .00 | -.02 | .13 |
| Green Odd 2 | — | -.06 | .35* | -.08 | 022 |
| Red Odd 1 | | — | .14 | .34* | -.09 |
| Red Odd 2 | | | — | .02 | -.11 |
| Red Odd Rule | | | | — | -.08 |
| **Laplacian Transformed** | | | | | |
| Green Odd 1 | .52* | -.07 | -.15 | -.19 | .44* |
| Green Odd 2 | — | .08 | -.01 | -.15 | .40* |
| Red Odd 1 | | — | .40* | .26 | .04 |
| Red Odd 2 | | | — | .51* | -.44* |
| Red Odd Rule | | | | — | -.34* |

*Note:* Correlations are reported controlling for variance due to age, sex, and minimum trial numbers. Green Odd 1 = ERP difference wave calculated from red standards minus green deviants in the first position of the stimulus pairs in the simple/green block, etc. “Odd” always refers to the color of the deviant stimulus in the relevant condition/stimulus position. Partial correlations were pooled across all imputed datasets. Correlations that are significant in the original dataset are marked with an asterisk (* = $p$ < .05; uncorrected for six experimental conditions).

**Supplementary Table S5**

Pooled partial correlations for ERP difference waves (vMMN) latencies

| Variable | Green Odd 2 | Red Odd 1 | Red Odd 2 | Red Odd Rule | Green Odd Rule |
|---|---|---|---|---|---|
| **Scalp Referenced** | | | | | |
| Green Odd 1 | .22 | .06 | -.14 | .14 | .09 |
| Green Odd 2 | — | -.20 | -.06 | -.06 | .29 |
| Red Odd 1 | | — | .44* | .06 | .07 |
| Red Odd 2 | | | — | .05 | .01 |
| Red Odd Rule | | | | — | -.03 |
| **Laplacian Transformed** | | | | | |
| Green Odd 1 | .26 | .04 | -.03 | .07 | .09 |
| Green Odd 2 | — | .33 | .07 | .07 | .28 |
| Red Odd 1 | | — | .29 | .40* | -.03 |
| Red Odd 2 | | | — | .50* | -.32 |
| Red Odd Rule | | | | — | -.18 |

*Note:* Correlations are reported controlling for variance due to age, sex, and minimum trial numbers. Green Odd 1 = ERP difference wave calculated from red standards minus green deviants in the first position of the stimulus pairs in the simple/green block, etc. “Odd” always refers to the color of the deviant stimulus in the relevant condition/stimulus position. Partial correlations were pooled across all imputed datasets. Correlations that are significant in the original dataset are marked with an asterisk (* = $p < .05$; uncorrected for six experimental conditions).

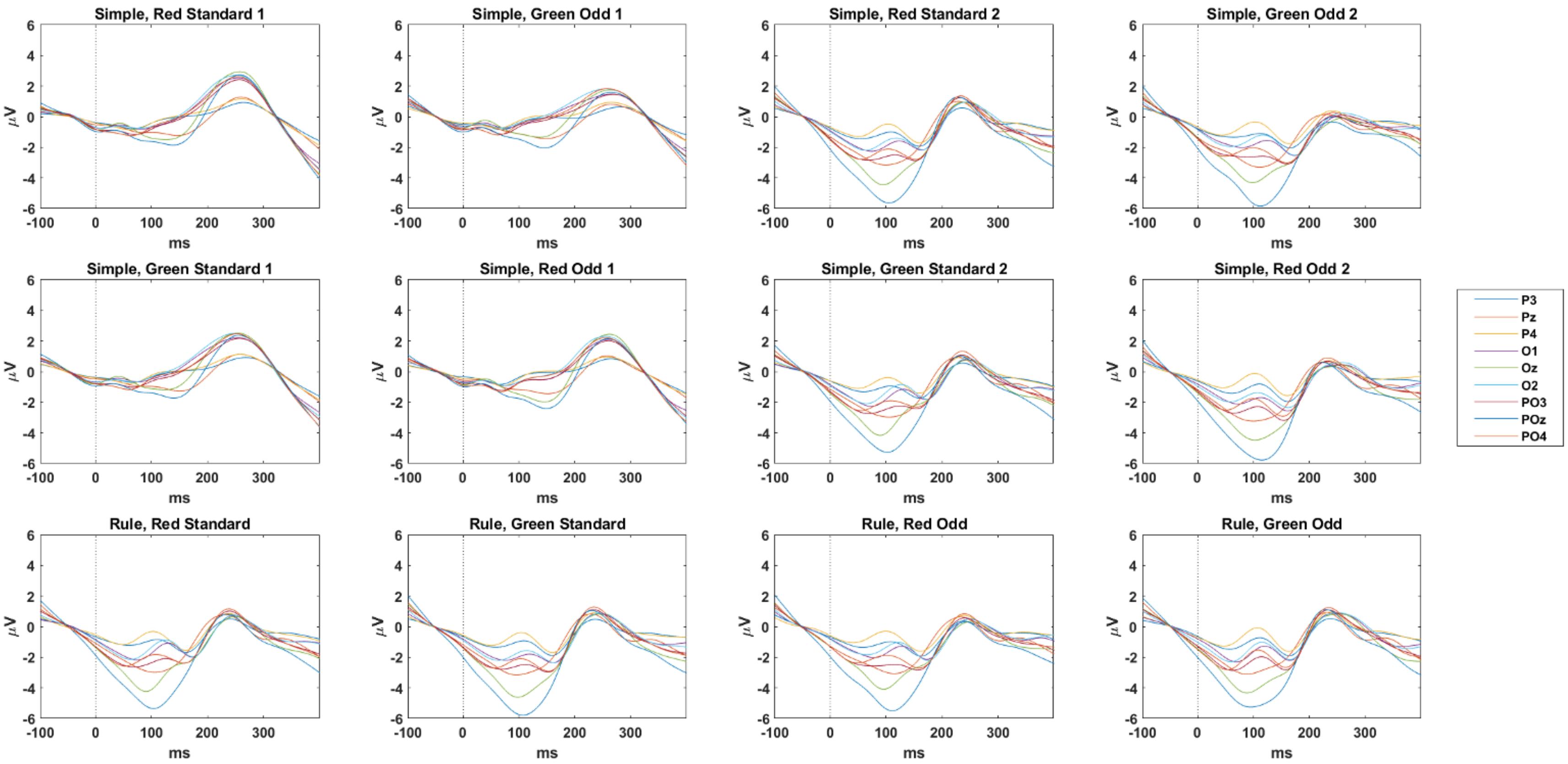


**Supplementary Fig. S1.** Plots depict the group-averaged, scalp-averaged reference ERPs for each stimulus condition, over all participants, in the nine electrodes of interest for that referencing approach. Variables extracted from these ERPs were used as inputs for the analyses of condition effects. Difference waves derived from these ERPs provided the basis for the analyses of individual differences effects (see Methods). Dotted lines indicate the time point of stimulus onset. Note the presence of anticipatory potentials in the pre-stimulus period for ERPs time-locked to the onset of second-position stimuli.

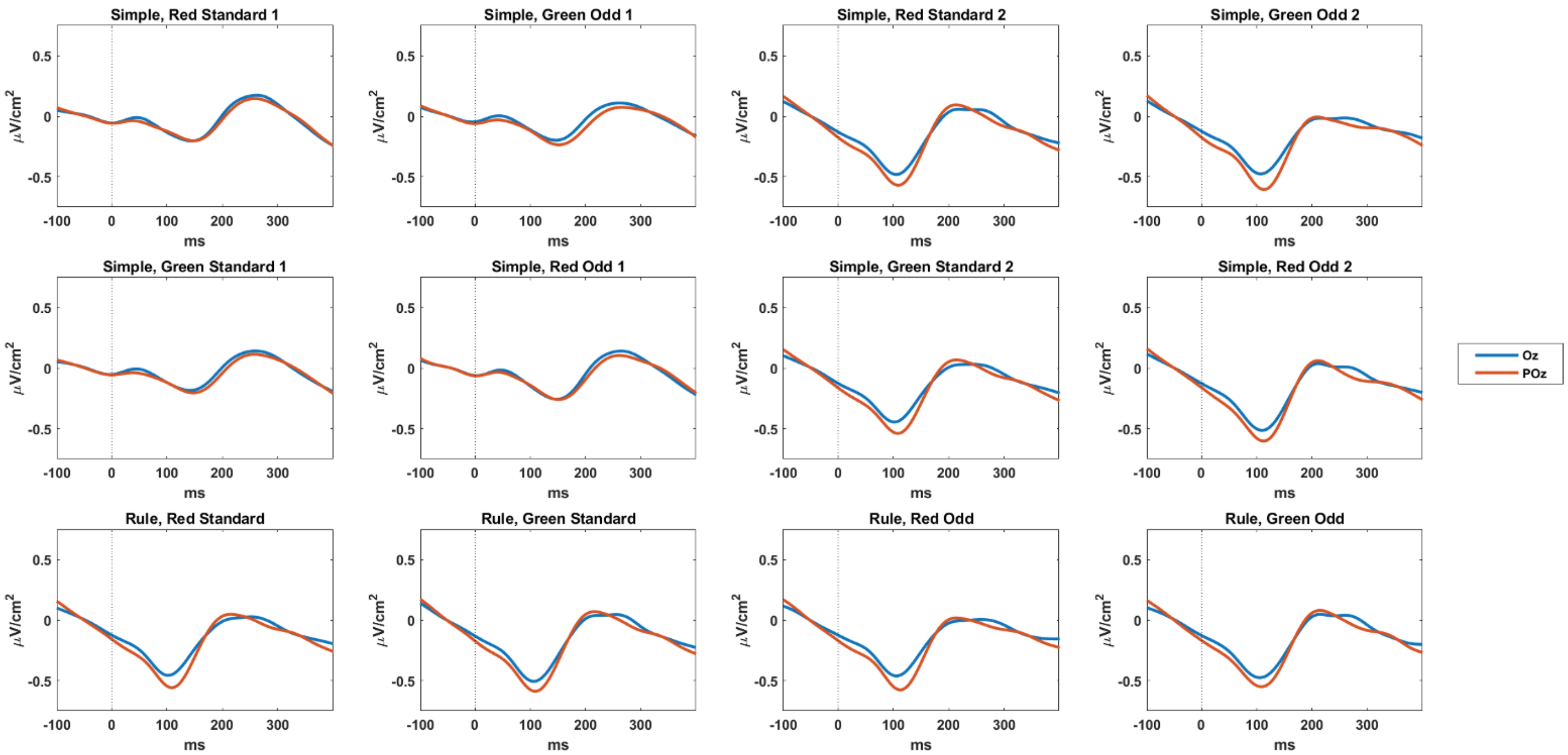


**Supplementary Fig. S2.** Plots depict the group-averaged, Laplacian transformed ERPs for each stimulus condition, over all participants, in the nine electrodes of interest for that referencing approach. Variables extracted from these ERPs were used as inputs for the analyses of condition effects. Difference waves derived from these ERPs provided the basis for the analyses of individual differences effects (see Methods). Dotted lines indicate the time point of stimulus onset. Note the presence of anticipatory potentials in the pre-stimulus period for ERPs time-locked to the onset of second-position stimuli.

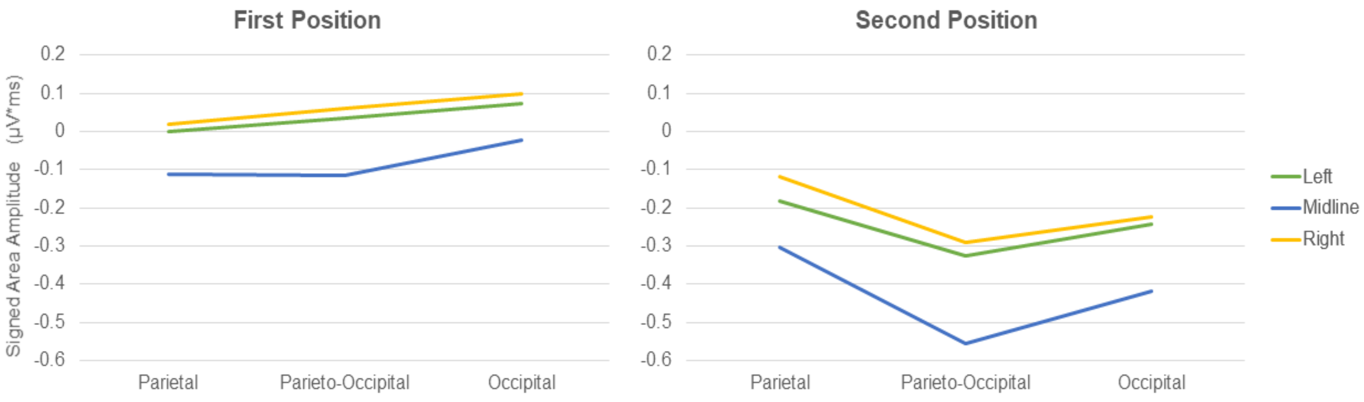


**Supplementary Fig. S3.** Interaction between stimulus Position, Anteriority, Hemisphere on the basis of group-averaged ERP amplitudes in the simple conditions. Values reflect negative integral ERP amplitudes for first and second-position stimuli, averaging over stimulus Deviance.

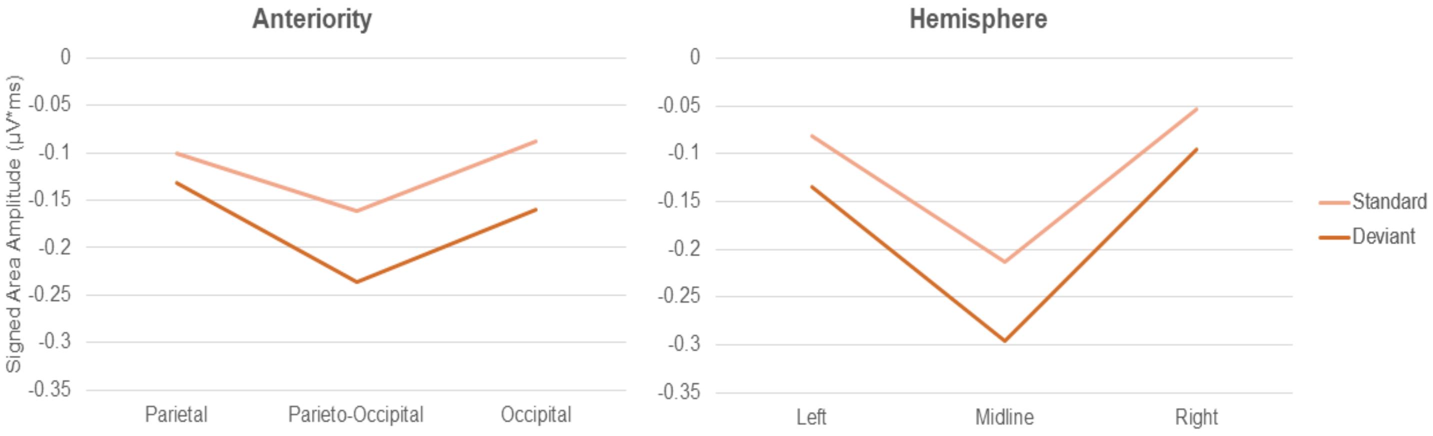


**Supplementary Fig. S4.** Interaction between stimulus Deviance and Anteriority and Deviance and Hemisphere on the basis of group-averaged ERP amplitudes in the simple conditions.

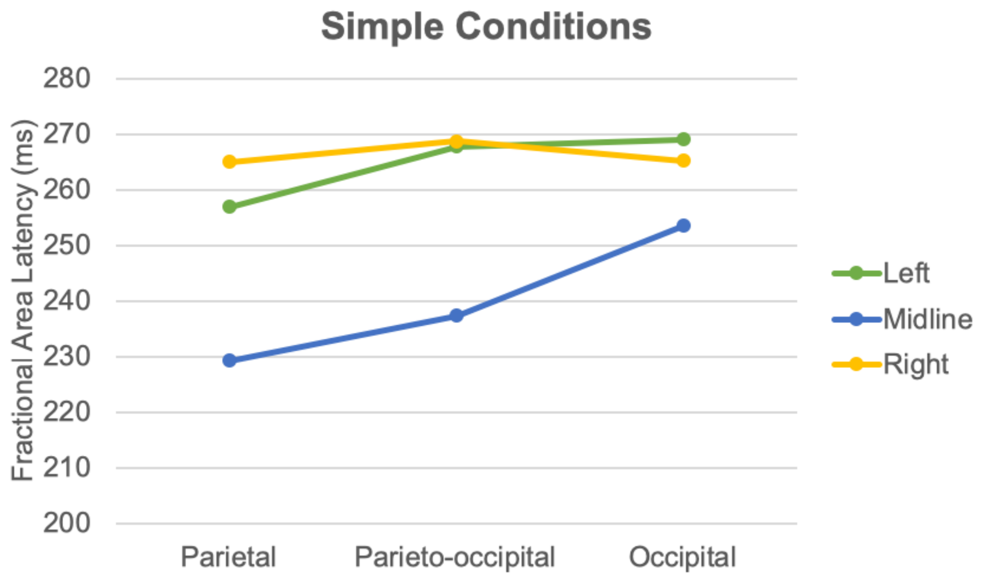


**Supplementary Fig. S5.** Interaction of Hemisphere and Anteriority on the basis of group-averaged ERP latencies in the simple conditions.

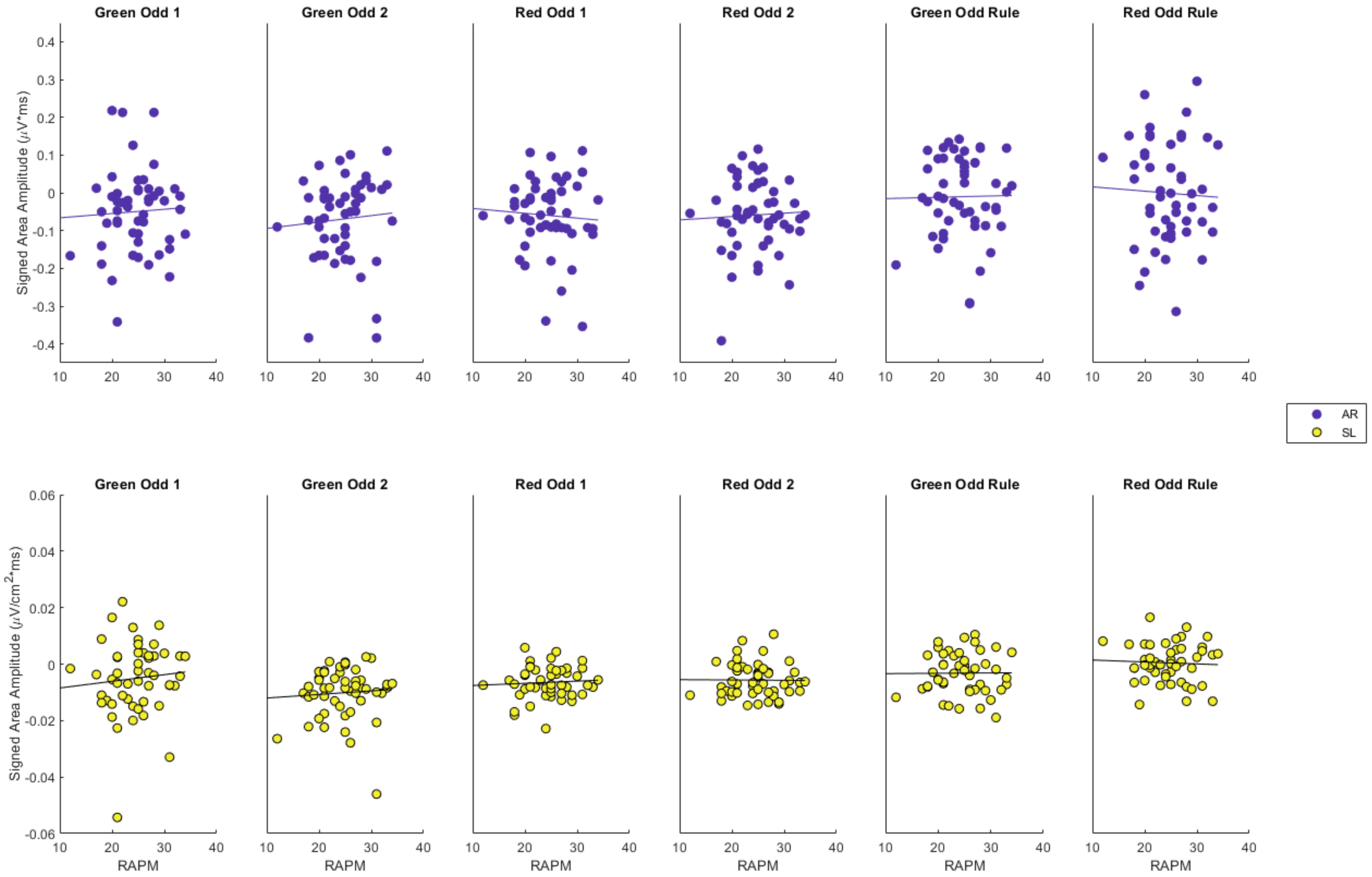


**Supplementary Fig. S6.** Scatterplots depict signed area amplitudes for each of the six vMMN difference waves plotted against RAPM scores, for the scalp-averaged reference and Laplacian-transformed data. Regression lines are plotted uncorrected for age, sex, and minimum trial numbers. RAPM = Ravens Advanced Progressive Matrices; AR = Scalp-averaged reference, SL = surface Laplacian.

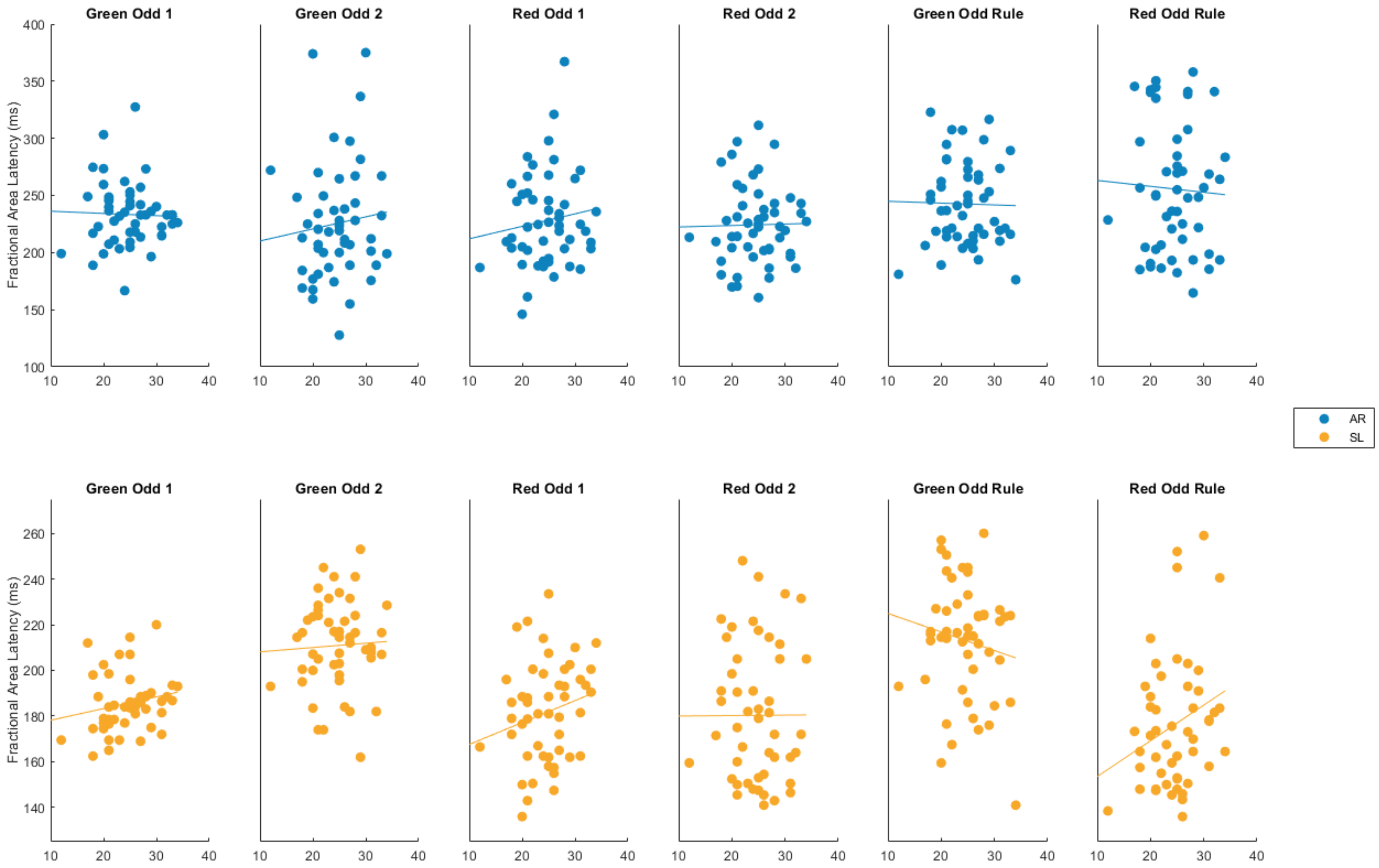


**Supplementary Fig. S7.** Scatterplots depict fractional area latencies for each of the six vMMN difference waves plotted against RAPM scores, for the scalp-averaged reference and Laplacian-transformed data. Regression lines are plotted uncorrected for age, sex, and minimum trial numbers. RAPM = Ravens Advanced Progressive Matrices; AR = Scalp-averaged reference, SL = surface Laplacian.
